# A bacterial cell wall repair and modification system to resist host antibacterial factors

**DOI:** 10.1101/2024.11.08.622053

**Authors:** Jessica Burnier, Clement Gallay, Kevin Bruce, Elisabet Bjånes, Louise Martin, Kinki Jim, Ho-Ching Tiffany Tsui, Amelieke Cremers, Johann Mignolet, Daniela Vollmer, Jacob Biboy, Victor Nizet, Waldemar Vollmer, Malcolm E. Winkler, Jan-Willem Veening

## Abstract

Pathogenic bacteria have acquired the ability to resist antibacterial defense mechanisms of the host. Streptococci are common in animal microbiota and include opportunistic pathogens like Group A Streptococcus (GAS) and *Streptococcus pneumoniae* (pneumococcus). While the conserved streptococcal S protein has been identified as a key factor in GAS virulence, its exact function is unclear. Here, we show that the pneumococcal S protein is crucial for resisting against host-derived antimicrobials by coordinating cell wall modification and repair. Specifically, we show that S proteins are septally localized through their transmembrane domain and contain an extracellular peptidoglycan (PG) binding LysM domain which is required for its function. Protein-protein and genetic interaction studies demonstrate that the pneumococcal S protein directly interacts with a PG synthase, class A penicillin binding protein PBP1a, and the PG deacetylase PgdA. Single-molecule experiments reveal that the fraction of circumferentially moving PBP1a molecules is reduced in the absence of S protein. Consistent with an impaired PBP1a function, streptococci lacking S protein exhibit increased susceptibility to cell wall targeting antibiotics and altered cell morphologies. PG analysis showed reduced N-deacetylation of glycans in the *S. pneumoniae* S protein mutant, indicating reduced PgdA activity. We show that pneumococci lacking the S protein cannot persist transient penicillin treatment, are more susceptible to the human antimicrobial peptide LL-37 and to lysozyme, and show decreased virulence in zebrafish and mice. Our data support a model in which S proteins regulate PBP1a activity and play a key role in coordinating PG repair and modification. This cell wall ‘sentinel’ control system provides defense against host-derived and environmental antimicrobial attack.

## Introduction

Streptococci are Gram-positive bacteria with a spherical or ovoid shape, often found as commensals in animals and humans. Notable species include *S. salivarius*, part of the human salivary microbiome^1^, and *S. thermophilus*, used in cheese production^2,3^. While typically harmless, some streptococci, such as *S. pyogenes* (group A *Streptococcus*, GAS) and *S. agalactiae* (group B *Streptococcus*, GBS), can cause serious diseases. GAS is responsible for approximately 700 million non-invasive infections each year (e.g., strep throat)^4–6^ and over 150,000 deaths from invasive conditions like necrotizing fasciitis and streptococcal toxic shock syndrome^5,7^. GBS significantly increases the chances of neonatal death by causing intra-amniotic and peripartum infections^8^. Another globally important human pathogen, *S. pneumoniae* causes nearly 100 million cases of lower respiratory infections and over half a million deaths each year^9^.

Streptococci utilize a variety of cell wall polysaccharides to adhere to their host and evade the immune system, enabling them to thrive as both commensals and pathogens. For instance, GAS and GBS possess unique rhamnose-glucose polysaccharides covalently attached to the cell wall peptidoglycan (PG) that play important roles in virulence^3,10^. Pneumococci employ PG-attached wall teichoic acids and exopolysaccharide capsules to avoid opsonization by immune cells^11–13^. In addition to these polysaccharides, species-specific surface proteins bound to PG, such as CbpA in *S. pneumoniae*, which interacts with human factor H^12^, and the GAS M protein, which recruits specific human proteins to resist immune clearance, are key to host adaptation^14^. These diverse cell surface components allow streptococci to adapt to various niches and lifestyles.

In addition to species-specific adaptations, all streptococcal genomes contain the *ess* gene, which encodes a highly conserved surface-exposed protein known as the S protein (**Fig. 1A, S1A**)^15^. The S protein is a critical determinant of immune evasion in GAS and GBS, with *ess* mutants exhibiting various phenotypes, including reduced binding to red blood cells and attenuated virulence^15,16^. For instance, the GAS S protein is believed to use red blood cells as a form of immune camouflage^15^. In GBS, absence of *ess* results in altered cell-surface properties and reduced levels of exopolysaccharides^16^. Despite these pleiotropic phenotypes on both the bacteria and host, the precise function of the streptococcal S protein has been unclear. Here, we show that the pneumococcal S protein (also known as SPV1429, SPD1429, SPR1457, or SP1604) is a central component of a PG repair and modification complex, including PBP1a and PgdA, which is necessary for protection against host innate defenses.

**Fig. 1:**
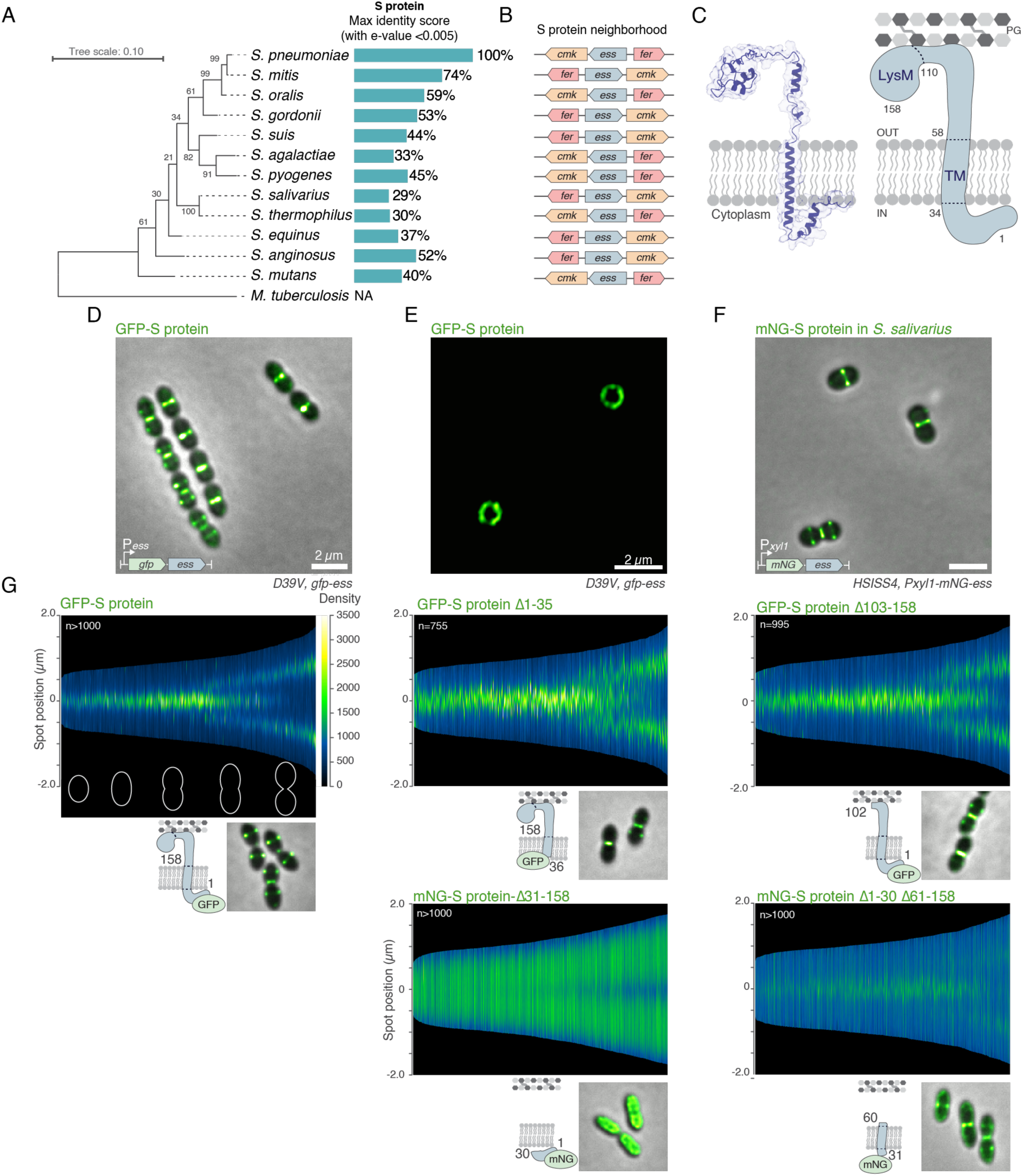
The S protein is localized at midcell through its transmembrane-spanning domain. (**A**) S protein conservation in Streptococci. Lineage tree based on 16S rRNA sequence of each species (see methods) was constructed using MEGA^36^. *Mycobacterium tuberculosis* was used as an outgroup and numbers represent bootstrap. Percentages indicate the percent identity of S protein for each species compared to *S. pneumoniae* S protein. (**B**) Gene co-occurrence neighborhood (data obtained from genome annotation in NCBI: see Methods). (**C**) AlphaFold model of the pneumococcal S protein. Cartoon of S protein with its predicted topology in the membrane and LysM domain binding to peptidoglycan (PG). Note that for clarity the ‘periplasmic space’ is drawn larger than the likely size^37^. (**D**) Deconvolved epifluorescence microscopy of live *S. pneumoniae* cells tagged with msfGFP-S protein that localizes at midcell. Scale bar = 2 μm. (**E**) 2D-SIM of msfGFP-S protein cells placed vertically in a microhole agarose pad. Scale bar = 2 μm. (**F**) Deconvolved epifluorescence microscopy of live *S. salivarius* cells tagged with mNeonGreen-S protein that localizes at midcell. Scale bar = 2 μm. (**G**) Localization signal of truncation fragments of S protein tagged with GFP or mNeonGreen (mNG) in approx. 1,000 *S. pneumoniae* cells, ordered by cell length and represented by heatmaps. Representative deconvolved microscopy images are shown (see **Fig. S3A** for more cells). Note that the schematics are not drawn to scale.

*S. pneumoniae* PBP1a is a class A penicillin binding protein (aPBP) involved in cell elongation, cell division and PG repair^11,17^. aPBPs are bifunctional enzymes that catalyze PG glycosyltransferase and transpeptidation, with PBP1a primarily localizing to the midcell region^18^. While PBP1a was initially found to be associated with the elongasome—a PG biosynthetic complex responsible for cell elongation^19–23—PBP1a^ may also play a significant role in PG repair, although the mechanisms underlying the regulation of the different roles of PBP1a in PG fortification and repair are not well understood^24,25^. Recent work in *E. coli* has similarly suggested that aPBPs are crucial for repairing the PG upon cell wall damage^26,27^. PgdA is an N-acetylglucosamine deacetylase that modifies glycan strands in PG by removing acetyl residues from GlcNAc residues^28^, thereby enhancing the bacterial cell wall’s resistance to lysozyme, an important component of the human innate immune system on mucosal surfaces^29^. A *pgdA* mutant showed reduced virulence in a murine intraperitoneal infection model^30^. Although the *pbp1a* and *pgdA* genes are not essential for *in vitro* growth, homologs of PBP1a and PgdA are found in all streptococci (**Fig. S2**), highlighting their importance in helping these bacteria resist and evade lysozyme and other host immune defenses^31^.

Using a combination of genetic, biochemical, cell biological, single-molecule, growth, and infection experiments, we demonstrate that the pneumococcal S protein colocalizes with PBP1a and PgdA. This triad plays a key role in defending against host-derived antimicrobials such as lysozyme and the cationic human cathelicidin antimicrobial peptide, LL-37. Phylogenetic analysis indicates that S proteins, along with their structural and functional homologues, are present not only in streptococci but in other Gram-positive bacteria such as *Bacillus subtilis* and *Listeria monocytogenes* (**Fig. S1**). We propose a model in which S proteins regulate the activity of PBP1a, and thereby PgdA, facilitating the repair of host-induced cell wall damage and timely deacetylation of newly synthesized PG, preventing lethal cell wall hydrolysis.

## Results

### Conserved midcell localization of the S protein

Phylogenetic analysis of the pneumococcal S protein (encoded by *spv_1429*) among streptococci showed a clear separation between S proteins from the mitis group and viridans streptococci, with *S. salivarius* having the most divergent sequence (**Fig. 1A**). Nevertheless, the gene neighborhood of *ess* is highly conserved in all streptococci, flanked by the conserved genes *cmk* and *fer* encoding cytidylate kinase and ferredoxin, respectively^32^ (**Fig. 1B**). An Alphafold model^33^ of the pneumococcal S protein predicts a cytoplasmic N-terminal domain, a single transmembrane domain, and an extracellular lysin motif domain (LysM) (**Fig. 1C**). LysM domains are typically involved in PG-binding^34^, suggesting a role of the S protein in cell wall biology. To explore this, we fused the S proteins from *S. pneumoniae* and *S. salivarius* with green fluorescent proteins (GFP or mNeonGreen) at their N-termini. Both fusions localized to the midcell (**Figs. 1D, 1F**), reminiscent of proteins involved in division and PG synthesis. For higher resolution S protein localization, we fabricated micropillars for vertical imaging of immobilized cells^35^. Structured illumination microscopy (SIM) of vertical *S. pneumoniae* GFP-S cells in micropillars revealed ring-like structures at the cell center (**Fig. 1E**). Further analysis with GFP or mNeonGreen fusions to various S protein truncations showed that the LysM domain was not essential for midcell localization, but the TM domain was crucial (**Fig. 1G**). Thus, the S protein exhibits conserved midcell localization in streptococci, dependent on its TM domain.

### The S protein is part of a protein complex including PBP1a and PgdA

To identify potential interaction partners of the pneumococcal S protein, we used the full-length S protein tagged at its N-terminus with GFP for immunoprecipitation using GFP nanobodies, followed by mass spectrometry. This approach revealed several proteins involved in cell wall biology, with PBP1a and PgdA being notably enriched (**Table S1**). To confirm these interactions, we constructed split-luciferase fusions of the S protein tagged with the large moiety of luciferase (LgBit) at either its N- or C-terminus. Correspondingly, several proteins, including PBP1a and PgdA, were tagged with the small moiety of luciferase (SmBit) (**Fig. 2A and S4**. When these moieties are in close proximity, they form a functional luciferase that emits light in the presence of a substrate^38,39^ (**Fig. 2A**). We then grew double-tagged strains in microtiter plates with luciferase substrate and measured growth and bioluminescence. The results showed strong and reproducible interaction signals between N-terminally tagged S protein and N-terminally tagged PBP1a and PgdA, as well as between C-terminally tagged S protein and C-terminally tagged PBP1a and PgdA (**Fig. 2B**). The split-luc interaction signal was either absent or not as strong for other potential partners identified by the pull-down experiment (**Fig. S4**), therefore in this study, we focused on the functional relationship of S protein with PBP1a and PgdA.

**Fig. 2:**
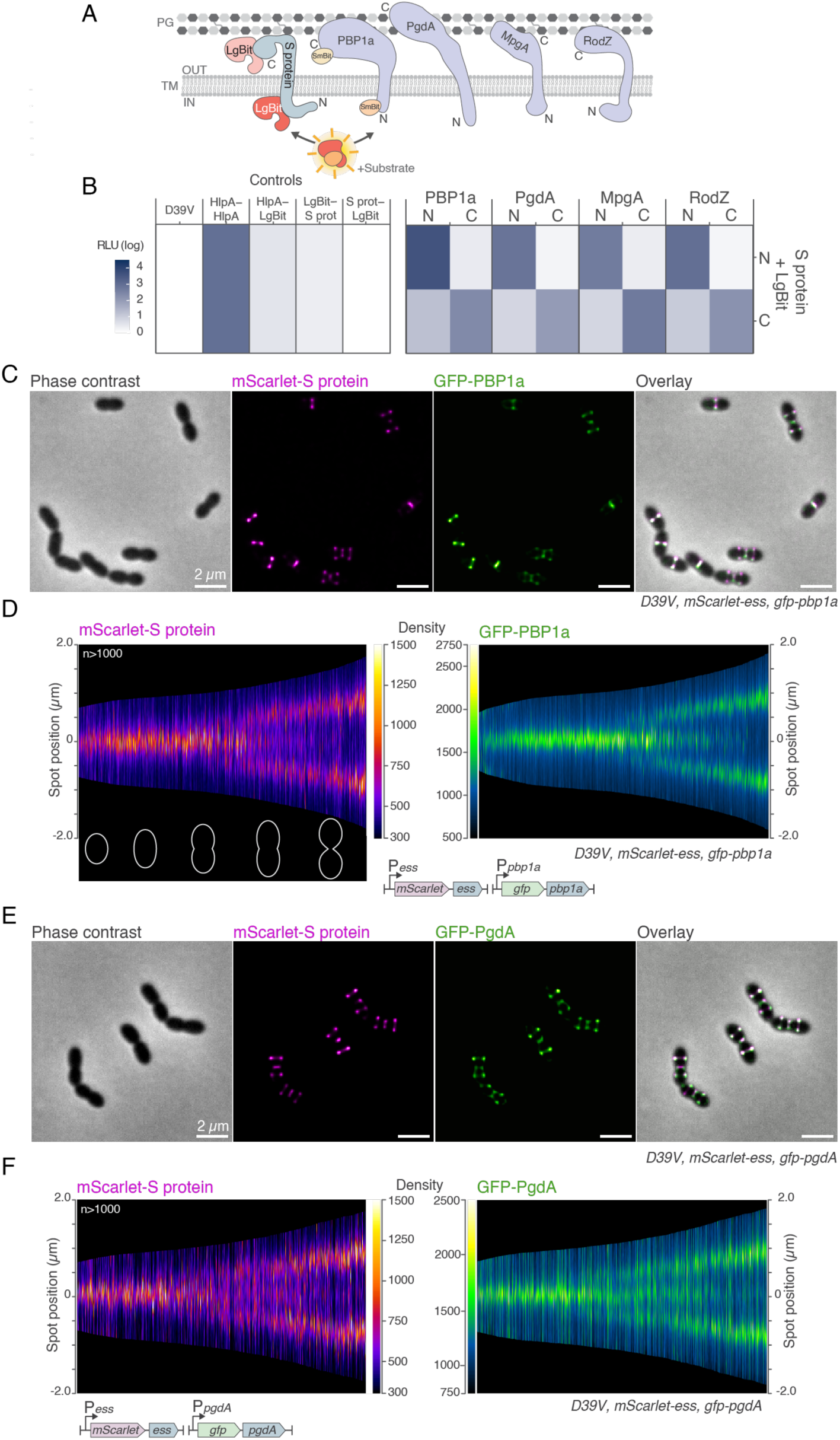
Pneumococcal S protein directly interacts with PBP1a and PgdA. (**A**) Schematic overview of detection of protein proximity through the split-luciferase assay. The S protein was fused to the LgBit moiety of a split luciferase at its N- or C-terminus and the targets to the SmBit part. For each interaction, four combinations of LgBit and SmBit associations were tested. (**B**) High relative luminescence unit (RLU, log) indicate close proximity. HlpA-LgBit/HlpA-SmBit (HlpA-HlpA) is used as a positive control^38,39^. C and N indicate the sub-cellular localization of the C- and N-terminal SmBit fusion, respectively. (**C-F**) mScarlet-S protein and msfGFP-PBP1a or msfGFP-PgdA co-localize. Double-labeled cells were grown in C+Y medium at 37°C and mid-exponentially growing cells were collected for fluorescence microscopy. Fluorescent proteins were expressed from their native locus as only copy in the cell. Scale bar = 2 µm. Fluorescent signal of one replicate from >1,000 cells per strain are ordered by cell length and represented by demographs plot performed using MicrobeJ^40^ (see Methods).

To further substantiate the interaction between the S protein, PBP1a, and PgdA, we constructed strains where the S protein was tagged with the red fluorescent mScarlet-I protein, and either PBP1a or PgdA was tagged with a monomeric super folder green fluorescent protein (GFP). While PBP1a midcell localization is well-established^18–20,24^, the localization of PgdA was previously uncharacterized. Our fluorescence microscopy analysis (**Fig. 2C-F)** revealed clear co-localization of the S protein with PBP1a and PgdA, with all three forming midcell rings. Furthermore, a cell cycle analysis of over 1,000 cells in exponential growth showed that the timing of midcell recruitment was identical for the three proteins (**Fig. 2D,F**). These findings support that the S protein directly interacts with PBP1a and PgdA, sharing membrane topology and co-localizing both spatially and temporally.

### The S protein is important for correct cell shape and tolerance to cell wall targeting antibiotics

Structural predictions and protein association studies show that the S protein is part of a complex minimally involving PBP1a and PgdA (**Fig. 2**). This suggests that the S protein, besides being a key virulence factor^15,16^, also plays a role in cell wall biology, since PBP1a is a PG synthase and PgdA a PG-modifying enzyme. To test this hypothesis, we created a Δ*ess* mutant in *S. pneumoniae* (Δ*ess^Sp^*) and examined growth, morphology and susceptibility to cell wall-targeting antibiotics. Cells were grown in microtiter plates in C+Y medium at 37°C (no added CO_2_), with optical density measured every 10 min. While the *S. pneumoniae* Δ*ess^Sp^* mutant showed similar growth to the wild type or a complemented mutant, increased lysis was observed in the late stationary phage (**Fig. 3A**). Most of this lysis was caused by the major autolysin LytA^41^, as shown by partial rescue in a *lytA* mutant background (**Fig. S3B**). Phase contrast and transmission electron microscopy (TEM) revealed that Δ*ess^Sp^* cells, overall, were slightly longer than wild type or complemented cells during the mid-exponential phase (**Fig. 3B-D**). However, throughout different growth phases, various morphological defects such as minicells, elongated cells, and chaining were observed (**Fig. S5**). The Δ*ess^Sp^*mutant also exhibited increased susceptibility to sub-MIC levels of beta-lactam antibiotics, including cefotaxime (CTX), piperacillin, and ceftriaxone (**Fig. 3E, Fig. S3C**), but not to the fluoroquinolone ciprofloxacin (**Fig. S3C**). Single cell analysis of CTX-treated cells showed significantly increased cell length (**Fig. 3F-G**). Interestingly, a truncated version of *ess* lacking the LysM domain also showed increased susceptibility to CTX, indicating that the LysM domain is required for S protein function (**Fig. 3H**).

**Fig. 3:**
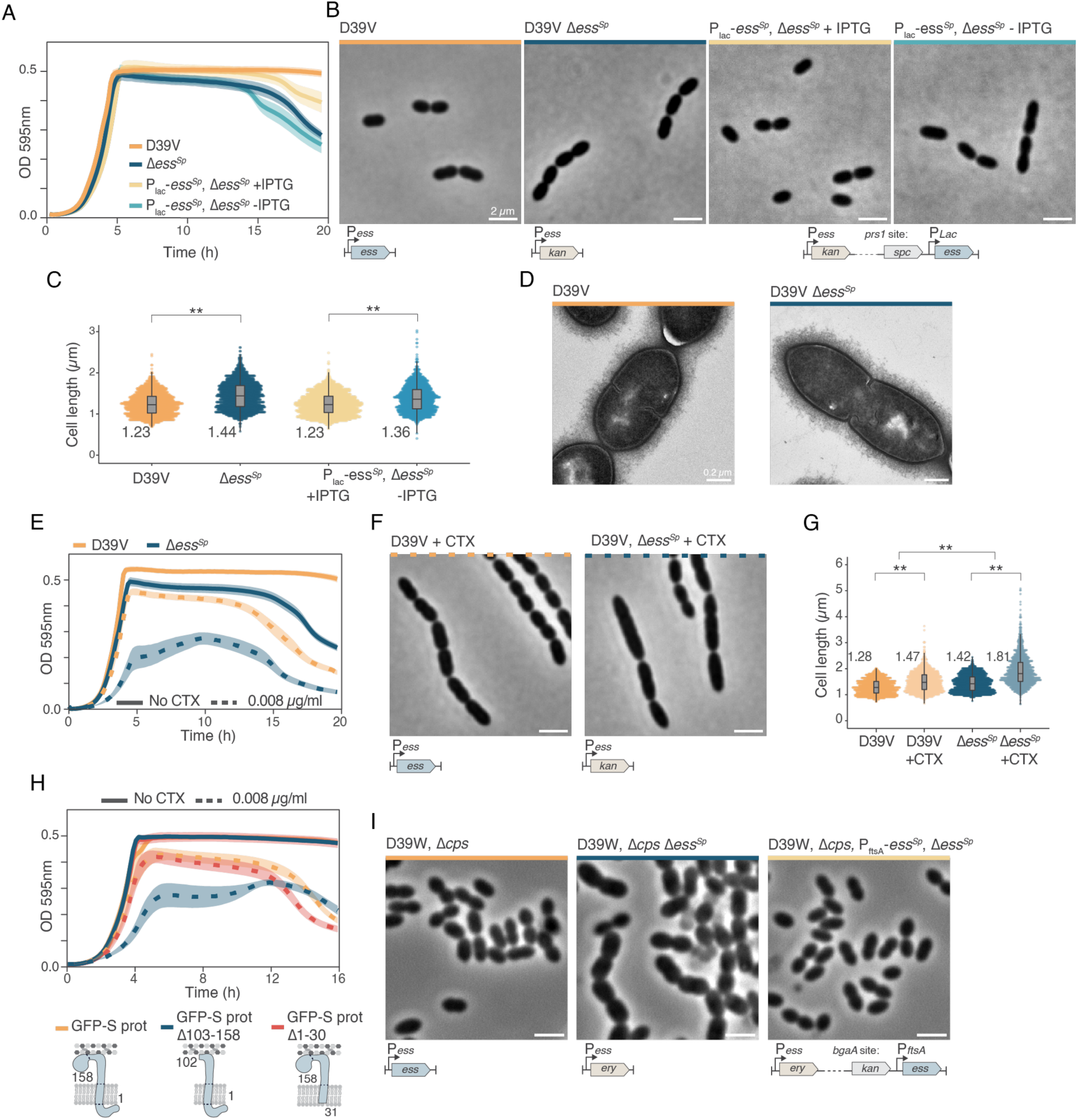
S protein is involved in cell wall homeostasis. (**A**) Growth curves of Δ*ess^Sp^* deleted pneumococcal cells shows increased cell lysis compared to D39V wild type at 37°C, which could be complemented by ectopic, IPTG inducible construct. For clarity, a linear scale for the optical density was used. Data are represented as the mean of n≥ 3 replicates and shading represents SEM. (**B**) Phase contrast microscopy of wild type and complemented Δ*ess^Sp^* strains in the encapsulated genetic background. Scale bar = 2 μm. (**C**) Quantitative analysis of phase contrast images show that Δ*ess^Sp^* cells are significantly longer than wild type or complemented cells. The number of quantified cells ranges between 1600 and 2,000 cells, numbers represent the means. ** p<0.01. (**D**) Transmission electron microscopy (TEM) images of D39V wild type strain and Δ*ess^Sp^*strain in the encapsulated genetic background. Scale bar = 0.2 μm. (**E**) Growth curves of pneumococcal cells subject to sublethal cefotaxime (CTX) at 37°C show that the Δ*ess^Sp^* mutant is more susceptible. Data are represented as the mean of n≥ 3 replicates and shading represents SEM. (**F**) Phase contrast microscopy images of *S. pneumoniae* grown in C+Y medium at 37°C and exposed to 0.008 μg/mL of cefotaxime (CTX). Scale bar = 2 μm. (**G**) Quantification of cell length of cells exposed to CTX (n>1600). See methods for statistical tests. (**H**) Growth curves of wild type and different truncated S protein variants treated with CTX (0.008 μg/mL). Data are represented as the mean of n≥ 3 replicates and shading represents SEM. (**I**) Phase contrast microscopy of D39W wild type, Δ*ess^Sp^* and complementation strains in the unencapsulated (Δ*cps*) genetic background. Scale bar = 2 μm.

The capsule is known to mask phenotypic effects of mutants involved in cell wall biology in *S. pneumoniae* ^18,42,43^. To further explore this, we generated Δ*ess^Sp^* mutants in a Δ*cps* genetic background. In this capsule-deficient context, the phenotypes of the Δ*ess^Sp^* mutants were exacerbated, displaying a variety of cell morphologies, including large, elongated cells and malformed minicells (**Fig. 3I & S6A**). These results suggest that streptococcal S proteins play a role in maintaining cell wall homeostasis.

### Genetic interaction analyses place the S protein as an integral partner of a cell wall repair and modification complex

To gain further insights into the function of the pneumococcal S protein, we performed a genome-wide synthetic lethal screen using CRISPRi-seq^44^ in the Δ*ess^Sp^* mutant background (**Fig. 4A**). We assessed the fitness of every gene in the *S. pneumoniae* D39V wild type genome and the Δ*ess^Sp^* mutant background by growing the CRISPRi libraries with and without IPTG to induce dCas9. The resulting fitness scores for each gene/operon were compared between the wild type and Δ*ess^Sp^* strains. This screen suggested that the S protein is involved in processes such as division, elongation, and cell wall biosynthesis, since genes known to be involved in these processes, such as *murE*, *divIB*, *rodZ*, and *mreD*^19,20,45,46^ became more essential in the absence of *ess^Sp^* (**Fig. 4A, Table S2**). Further investigations confirmed that an *ess*/*rodZ* double mutant could not be constructed (**Fig. 4B)**. In line with previous findings that a Δ*pbp1a* deletion suppresses the requirement for RodZ^22^, whole genome sequencing (WGS) of a single *rodZ* mutant revealed a suppressor mutation in *pbp1a* (Y191C) (**Fig. 4B**). Notably, the deletion of *pbp1a* also alleviates the requirements of the otherwise essential muramidase MpgA^22^.

**Fig. 4:**
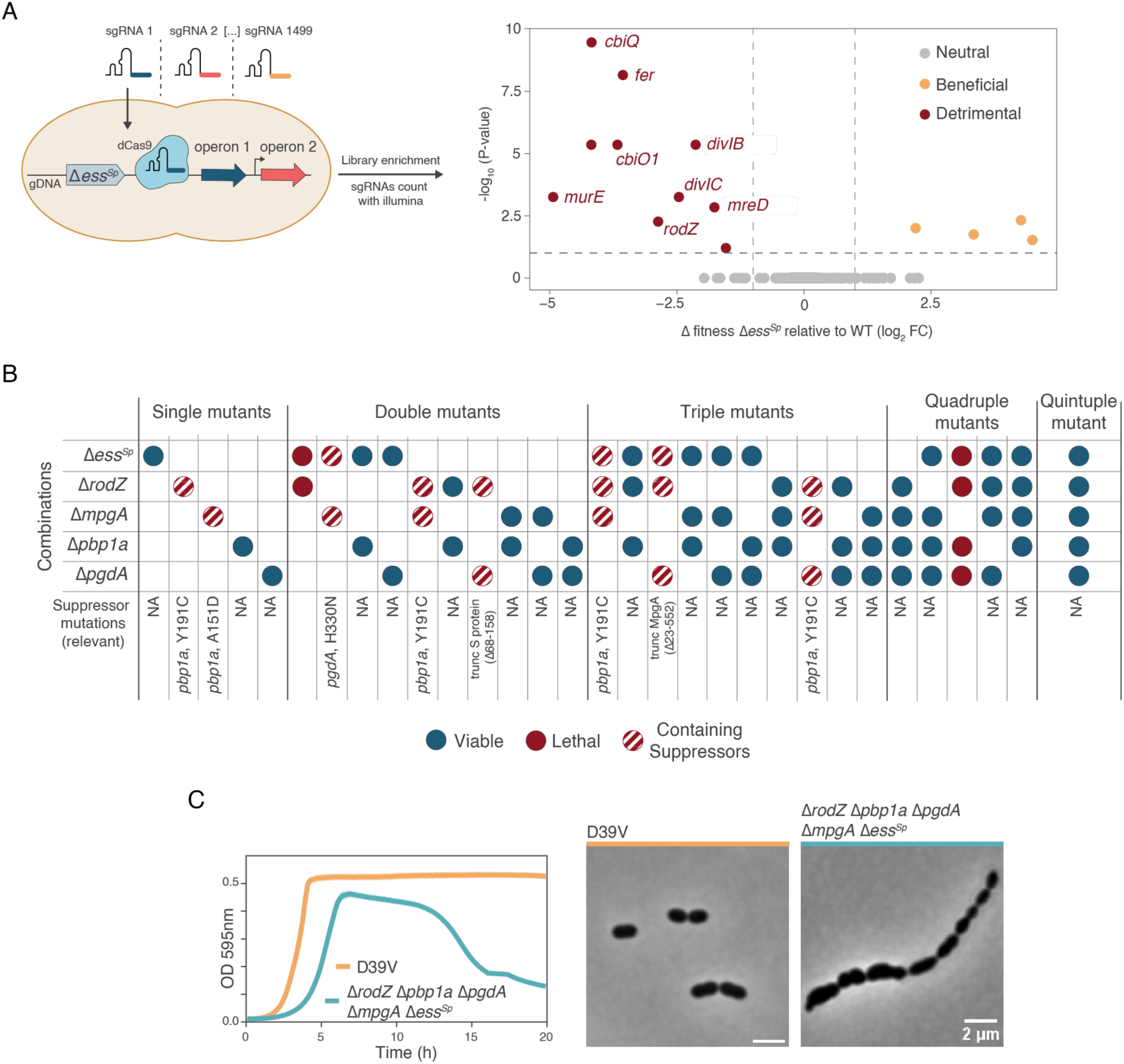
*ess^Sp^* is genetically linked to *pbp1a*, *pgda*, *mpgA* and *rodZ*. (**A**) A genome-wide sgRNA library was introduced in both wild type and an Δ*ess^Sp^*mutant background. Comparison of fitness cost of gene depletion by CRISPRi between the wild type background and Δ*ess^Sp^* mutant background is shown with putatively synthetic lethal interactions highlighted in red. (**B**) Genetic interactions with PgdA, PBP1a, MpgA, RodZ. Each column represents a strain with the corresponding genes deleted (dots represent deletions). Plain blue dots symbolize strains without relevant suppressors, dashed red dots symbolize strains containing a relevant suppressor, and red dots symbolize combinations of deletions that are lethal and could not be made. See **Table S3** for suppressor details of each strain. (**C**) Left: Growth curves of Δ*ess^Sp^,* Δ*pbp1a,* Δ*pgdA,* Δ*mpgA,* Δ*rodZ* deleted pneumococcal cells shows impacted growth compared to D39V wild type at 37°C. For clarity, a linear scale for the optical density was used. Data are represented as the mean of n≥ 3 replicates. Right: Phase contrast microscopy of wild type and Δ*ess^Sp^,* Δ*pbp1a,* Δ*pgdA,* Δ*mpgA,* Δ*rodZ* pneumococcal strain in the encapsulated genetic background. Scale bar = 2 μm.

Since the pneumococcal S protein directly interacts with PBP1a and PgdA, and MpgA and RodZ are known to demonstrate midcell localization patterns^24^, we investigated whether the S protein also interacts with RodZ and MpgA using our split-luc assay. Alphafold predictions indicated that both RodZ and MpgA share similar membrane topologies with the S protein (**Fig. 2A, S4A**). Our results showed that both N-terminally tagged and C-terminally tagged S proteins interacted with likewise tagged RodZ and MpgA (**Fig. 2B)**.

The results presented so far demonstrate that the S protein is part of a midcell localized complex that includes PBP1a, PgdA, RodZ, and MpgA. To explore genetic interactions within this network, we systematically generated knockout strains and combinations of deletion mutants, followed by WGS of the resulting strains (**Fig. S6-S9, Table S3**). As shown in **Fig. 4B**, this complex appears to be closely interconnected, as certain mutants and combinations could only be constructed when suppressor mutations arose in one of these five genes. For instance, a *rodZ*/*pgdA* double deletion mutant could not be readily obtained. However, performing WGS on a rare, large growing, suppressor colony containing the correct *rodZ*/*pgdA* double mutation, showed that this clone also acquired an inactivation mutation in *ess^Sp^*(*ess^Sp^* Δ68-158). Similarly, an *ess^Sp^*/*mpgA* double mutant was only viable with a suppressor mutation in *pgdA* (H330N). Interestingly, a quintuple mutant lacking Δ*ess^Sp^*, Δ*pbp1a*, Δ*pgdA*, Δ*rodZ*, and Δ*mpgA* was viable, without evidence for additional suppressor mutations, albeit with highly perturbed cell growth and morphology (**Fig. 4C)**. Thus, for cell viability, there is a crucial need for balanced enzymatic activities within this complex, suggesting that RodZ and the S protein, which lack predicted enzymatic functions, may serve as important adaptors that coordinate or regulate the activities of PBP1a, MpgA, and PgdA.

### Reduced fraction of circumferentially moving PBP1a in the absence of the S protein

The genetic interaction studies demonstrate that the S protein is part of a complex centered around PBP1a, which is known to be linked to cell elongation and cell wall repair^17,24,25^. To test whether the function of PBP1a is perturbed by the absence of the S protein, we performed single-molecule imaging of HaloTag-PBP1a. Our previous work confirmed that this fusion is functional and showed that a fraction of PBP1a molecules move circumferentially at midcell. Active PG synthesis was shown to drive circumferential midcell motion with velocities similar to those of core-elongasome components PBP2b, MreC, and RodA^24^. Another fraction of PBP1a molecules diffuse over the cell surface and exhibit short (<1 s) pauses, which may reflect PG damage repair^24^. As shown in **Fig. 5A**, in the absence of the S protein, a smaller fraction of PBP1a molecules moved circumferentially, a condition that could be complemented by an ectopic construct. Notably, the velocity and duration of circumferentially moving PBP1a did not change upon deletion of *ess* (**Fig. S10 A-C**). These results suggest that the Δ*ess^Sp^* phenotype may be partly due to the observed effects on PBP1a. Moreover, a Δ*pbp1a* mutant displayed increased susceptibility to CTX, and a Δ*pbp1a* Δ*ess^Sp^*double mutant did not exacerbate the Δ*ess^Sp^*phenotype (see arrow, **Fig. 5B**). Interestingly, a Δ*pgdA* mutant did not show reduced susceptibility to CTX (**Fig. S3D**), suggesting that a main function of the S protein may be to control PBP1a activity. Importantly, deleting *ess* did not affect the frequency or velocity of circumferentially moving molecules of iHT-PBP2b (**Fig. S10D-E**). The duration was reduced in the absence of the S protein, but both the WT and the mutant values (28 and 21 s, **Fig. S10E**) are similar to previous measurements of elongasome proteins^24^. Altogether, these results suggest that removal of the S protein significantly impairs PBP1a function.

**Fig. 5:**
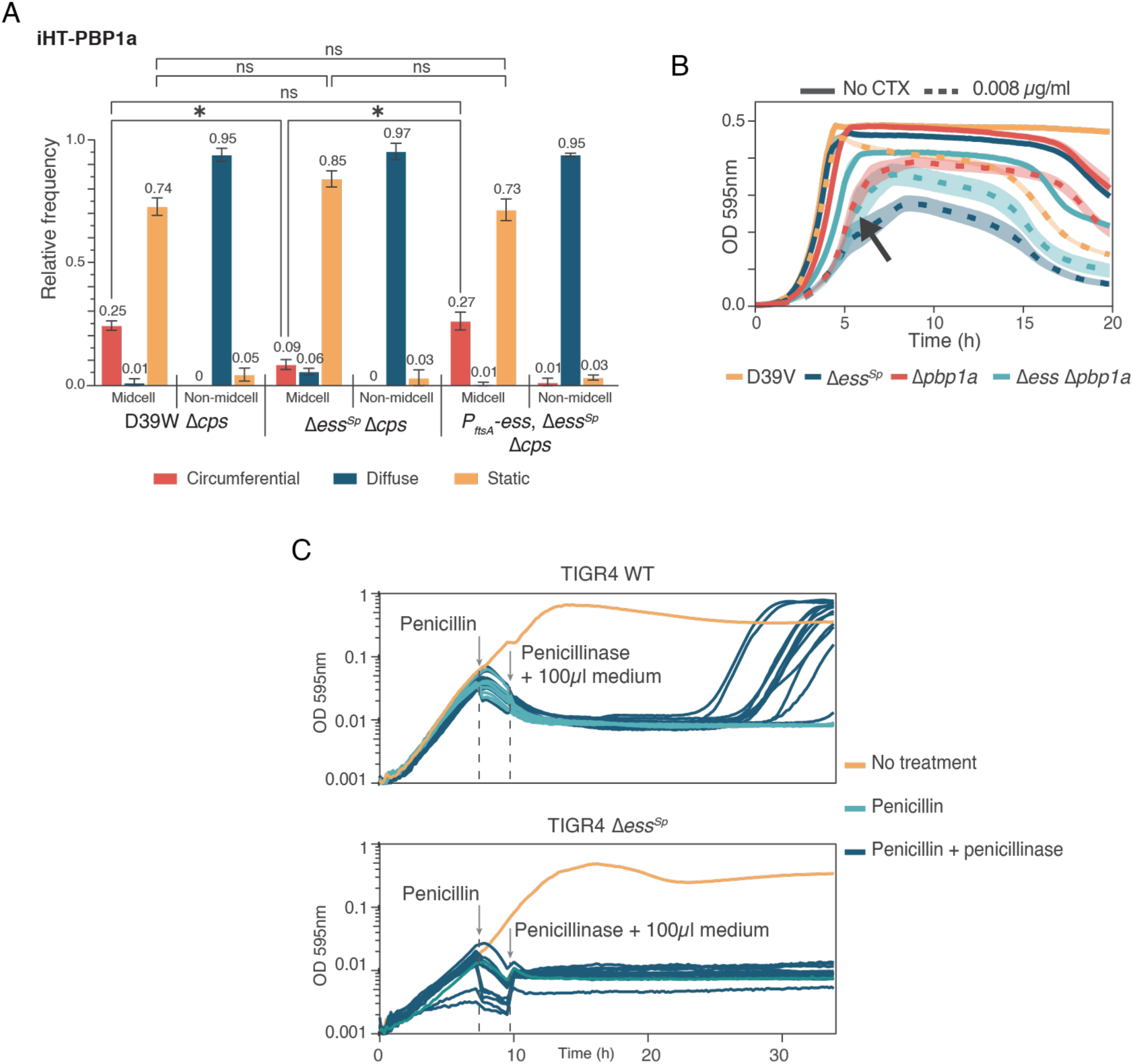
The effects on PBP1a contribute to Δ*ess^Sp^* phenotypes. (**A**) Deleting *ess* decreases the frequency of circumferentially moving iHT-aPBP1a in unencapsulated D39W. All strains express a functional fusion of the HaloTag (iHT) domain fused to PBP1a. Movement patterns of HT-labeled single molecules of PBP1a are shown. Bars represent the mean relative frequencies (±SD) of circumferential, diffusive and static molecules at midcell or away from the midcell (non-midcell) as determined by live-cell TIRFm. For each strain, ≥100 molecules were analyzed at both midcell and non-midcell from two biological replicates, with mean values shown above the bars. * P ≤ 0.05; ns (not significant) P > 0.05. See Methods (Single-molecule dynamics) for details of statistical tests. (**B**) Growth curves of pneumococcal cells subject to cefotaxime (CTX) at 37°C show that the Δ*ess^Sp^*, Δ*pbp1a* and Δ*ess^Sp^* Δ*pbp1a* mutants are more susceptible than the wild type strain. Data are represented as the mean of n≥ 3 replicates and shading represents SEM. (**C**) Penicillin persistence assay. Growth curves of pneumococcal serotype 4 strain (TIGR4) wild type and Δ*ess^Sp^.* Penicillin G was added to the media followed by the addition of 0.0125 units of penicillinase 2h later. Persistence was observed in the TIGR4 wild type strain, while the Δ*ess^Sp^* mutant was not able to persist antibiotic treatment. Abrupt changes in the curves are due to the plate reader being opened and to volume changes. Subsequently, TIGR4 WT strain penicillin susceptibility was confirmed using oxacillin disk diffusion method.

### The S protein is required for penicillin persistence

Since PBP1a is also implicated in penicillin tolerance and resistance^13^, this raises the question of whether a functional S protein is required for bacterial persistence^47^. To test this, we established a penicillin persistence assay where exponentially growing cells were exposed to a 100xMIC dose of penicillin G (PenG), followed by the addition of 0.0125 units of penicillinase 2h later (**Fig. 5C**). The penicillinase degrades PenG in the growth media, allowing surviving cells, potentially persister cells^48^, to regrow. Using this protocol, we did not observe any survival in *S. pneumoniae* strain D39V. However, in a serotype 4 strain (TIGR4), persistence was observed, with 11 out of 12 cultures surviving transient penicillin treatment. Strikingly, none of the cultures of a Δ*ess^Sp^* mutant in the TIGR4 background were able to persist through antibiotic treatment (**Fig. 5C**).

### Reduced activity of PgdA in Δ*ess^Sp^*

The results so far indicate that the cell envelope of *ess* mutants is weakened. To test whether the PG profile is altered in the Δ*ess^Sp^* mutant, we isolated cell walls and PG from exponentially growing cells and performed reverse-phase HPLC to quantify the muropeptides^49^. To facilitate analysis, all strains lacked a capsule^49^. We also included a Δ*pgdA* strain, which is known to lack deacetylated muropeptides, serving as a reference to identify and assign the peaks corresponding to different peptides^28,49^. As shown in **Fig. 6A**, the global PG profile of the Δ*ess^Sp^* mutant was largely similar to the wild type. However, there was a modest reduction in the amount of deacetylated muropeptides in the Δ*ess^Sp^* mutant, whereas overproduction of the S protein may have resulted in a slight increase in deacetylation (**Table S4**). Specifically, the proportion of deacetylated disaccharide tripeptide (peak 2) was 5.1% in the wild type, 5.7% in the strain overexpressing S protein, and 4.0% in the Δ*ess^Sp^* mutant. As expected, deacetylated muropeptides were absent in the Δ*pgdA* mutant (**Fig. 6A, Table S5**). These results are consistent with a small decrease in PgdA function in the absence of the S protein, possibly due to impaired synthesis of nascent PBP1a-derived PG, a possible substrate for PgdA (**Fig. 5A**).

**Fig. 6:**
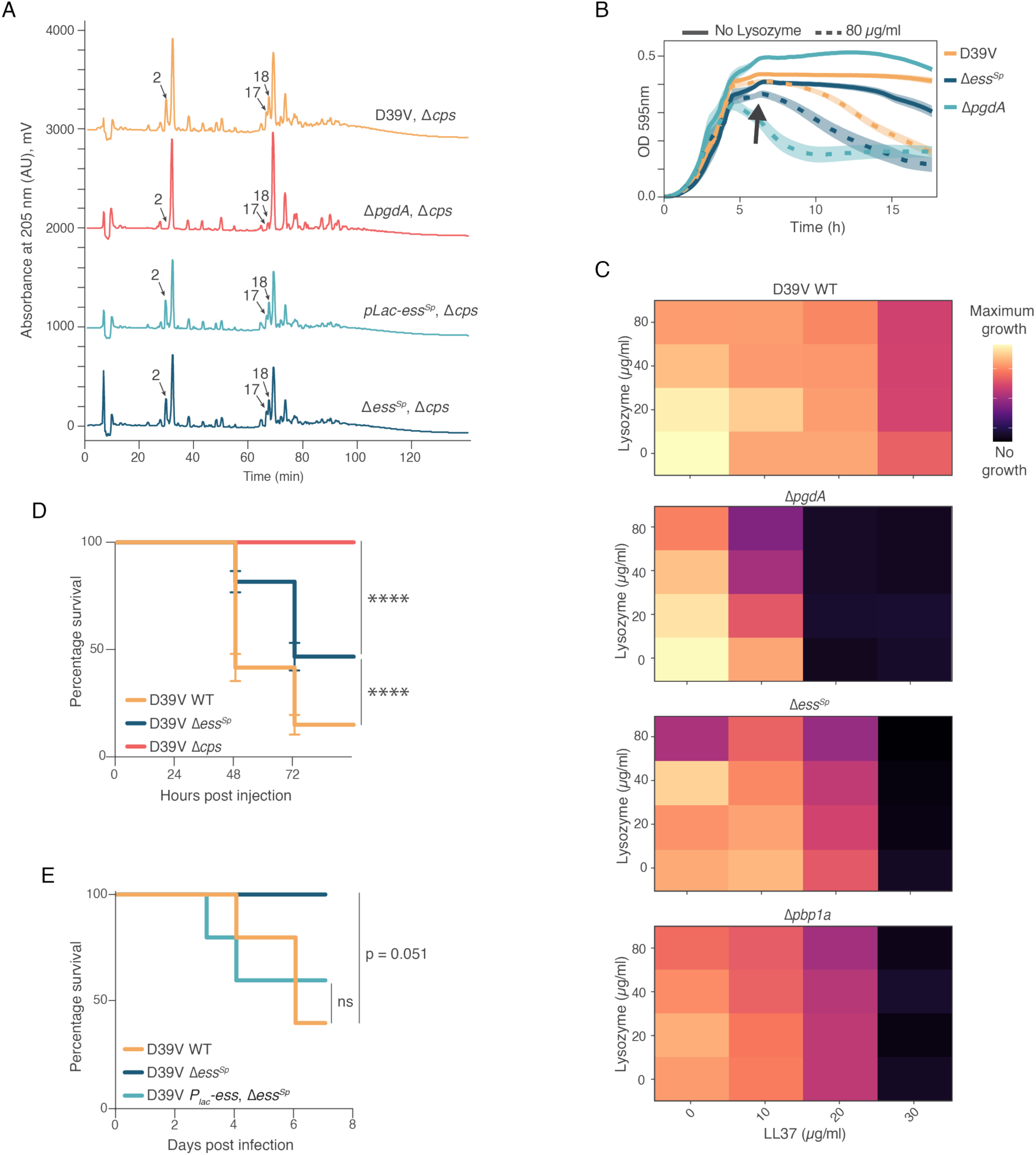
S protein is required to protect streptococcal cells from damages inflicted by the host innate immune system. (**A**) Muropeptide profiles of exponentially growing cells in C+Y medium at 37°C were obtained by reversed-phase HPLC^49^. The area under each peak (numbers) were calculated for each strain (**Table S5**). Peak number 2, 17 and 18 correspond to Tri[deAc], TetraTri[deAc]^‡^, and TetraTri[deAc]^‡^, respectively^49,53^. As reported before by (Bui et al. 2011). (**B**) Growth curves of cells subject to 80µg/mL lysozyme (dashed lines), show that Δ*ess^Sp^* mutant is more susceptible to lysozyme than wild type bacteria. Data are represented as the mean of n≥ 3 replicates and shading represents SEM. (**C**) Heatmap of the area under the curve (AUC) of the growth curves in liquid media. Lysozyme and LL-37 act synergistically on the Δ*ess^Sp^* mutant. Empirical AUC was plotted between 0-7 hours. (**D**) Survival curves of two days post fertilization zebrafish larvae injected with 200 CFUs of *S. pneumoniae* D39V wild type, Δ*ess^Sp^* mutant or Δ*cps* mutant into the hindbrain ventricle. The data represent the mean ± SEM of three biological replicates with 20 larvae per group (n = 60 in total/group); all groups are significantly different to each other; ∗∗∗∗ P value <0.0001 (Mantel-Cox test). (**E**) Mortality curves in B6 mice infected with 5*10^6^ CFUs of *S. pneumoniae* D39V wild type, Δ*ess^Sp^*mutant or P*_lac_-ess* Δ*ess^Sp^* complementation intratracheally. Notably, there is a constitutive ectopic expression of *ess* in the complementation strain as no *lacI* is present in the strain. Mice were monitored for survival for 7 days (n=5/group). P value = 0.051 (3/5 dead vs 0/5 dead), ns = not significant (p value = 0.814).

### The S protein is crucial for protection from host-induced cell damage

Since the deacetylation of PG by PgdA is required for high level resistance to human lysozyme^30,31^, we tested the susceptibility of the Δ*ess^Sp^*mutant. In line with the muropeptide analysis, the Δ*ess^Sp^* mutant was indeed more susceptible to lysozyme than wild type bacteria (see arrow, **Fig. 6B**). In summary, these results so far suggest that the phenotypes displayed in the Δ*ess^Sp^* mutant are partly due to disrupted activities of both PBP1a and PgdA, leading to increased susceptibility to autolysis, cell wall-targeting antibiotics, and lysozyme. Beside lysozyme, mammalian hosts produce antimicrobial peptides such as LL-37 (cathelicidin), a cationic peptide. Cationic antimicrobial peptides (CAMPs) are produced by epithelial cells and immune cells as part of the innate immune response to pathogens. CAMPs are amphipathic, cytotoxic, pore-forming peptides with strong antimicrobial activity^50,51^ that trigger a cell envelope stress response^52^. To test whether the S protein, PBP1a, and PgdA play roles in responding to host-induced damage, we performed checkerboard assays to assess the susceptibility of mutants towards both human LL-37 and lysozyme. As shown in **Fig. 6C**, LL-37 and lysozyme acted synergistically in all three mutants, which were substantially more inhibited than the wild-type. Importantly, the increased susceptibility of the S protein can be complemented (**Fig. S11**).

### Reduced virulence of the Δ*ess^Sp^* mutant

Finally, we tested whether *S. pneumoniae* lacking the S protein were more easily cleared by the host immune system using infection assays in a zebrafish meningitis model and a mouse pneumonia model. As shown in **Fig. 6D**, a higher survival rate was observed in zebrafish larvae challenged with the Δ*ess^Sp^* mutant compared to those challenged with the wild type. A similar trend was seen in the mouse model; all mice infected with the Δ*ess^Sp^* mutant survived, while mice challenged with wild type or complemented S protein bacteria succumbed to disease (**Fig. 6E**).

## Discussion

The original locus tag of the GAS S protein is SPy_0802. Due to its high conservation among GAS strains and homologous proteins in other streptococci, it was named the S protein^15^. S proteins of GAS and GBS were shown to be important for bacterial survival in various infection models^15,16^ and have emerged as potential vaccine candidates for preventing streptococcal infection^54^. Here, we show that the S protein is an integral component of a cell wall repair complex comprising PBP1a, PgdA, and possibly MpgA and RodZ (**Fig. 7, S12**). Our data support a model in which cell envelope damage— caused by antibiotics or host-derived antimicrobials such as lysozyme—is repaired by PBP1a and modified (through deacetylation) by PgdA to prevent further damage. In this scenario, fast-diffusing PBP1a molecules pause to repair damaged PG, while circumferentially moving PBP1a complexes are involved in PG fortification and possibly repair. The serine/threonine kinase StkP and adaptor protein GpsB are likely also involved in sensing this damage^19,55,56^. At the damaged site, the S protein activates PBP1a to produce nascent PG, which PgdA then further matures by deacetylation (**Fig. 7C**). This coordination of PBP1a and PgdA activity ensures timely PG restoration. The exact roles of MpgA and RodZ remain unclear, but they may help in releasing the PBP1a/PgdA complex from repaired sites, allowing it to address additional damage (**Fig. 7C**). In the absence of the S protein, the PBP1a/PgdA complex localizes to the damaged site but is not activated, and thus remains stuck and cannot efficiently repair and modify subsequent cell wall damage (**Fig. 7C**). This leads to a reduced fraction of circumferentially moving PBP1a molecules, accumulation of cell wall defects and enhanced sensitivity to cell wall attack in the Δ*ess^Sp^* mutant.

**Fig. 7:**
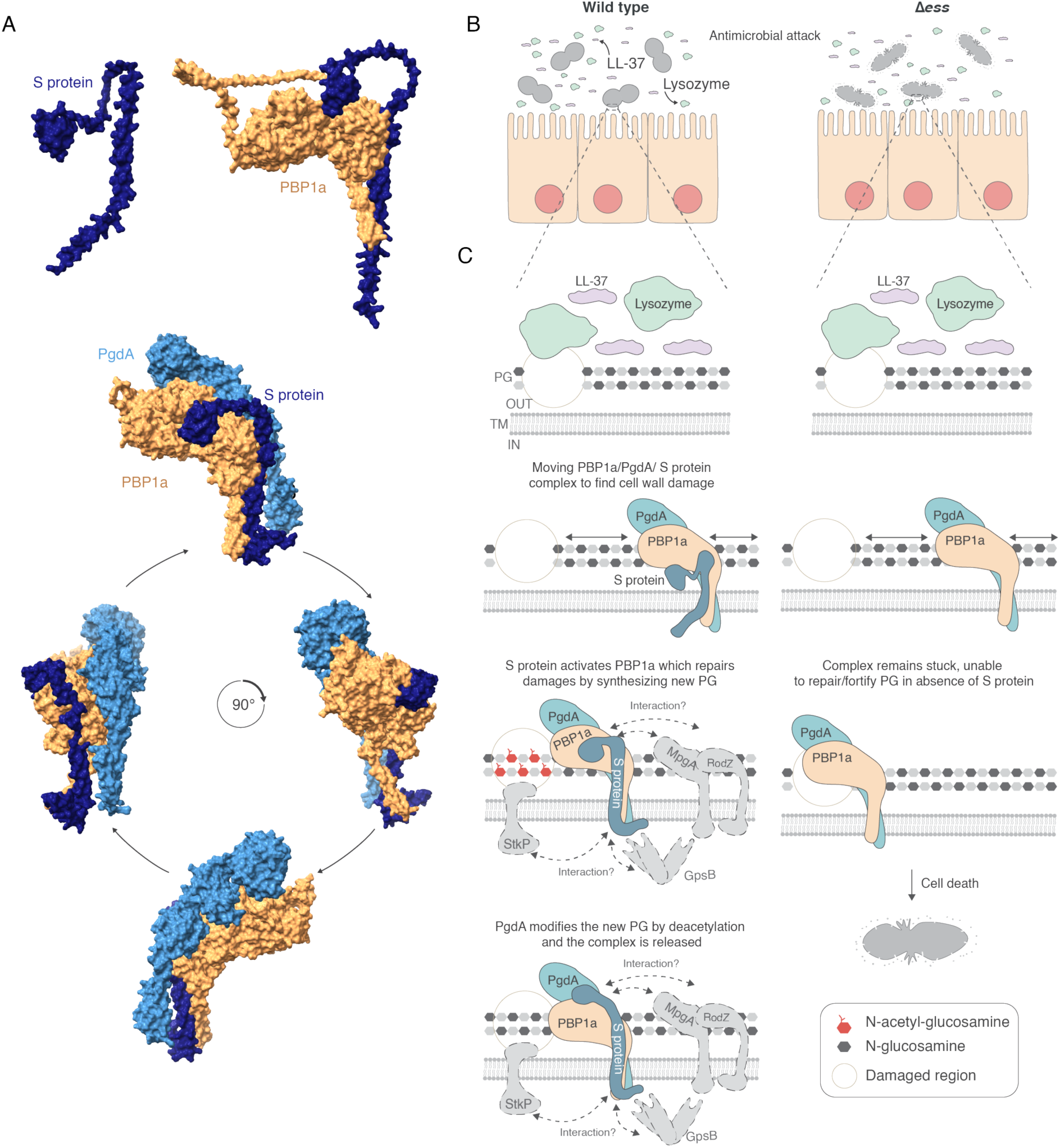
Streptococcal S protein coordinates the PBP1a:PgdA sentinel cell wall repair-modification complex required for protection from host antimicrobial challenges. (**A**) Alphafold structural predictions of the S protein in complex with PBP1a and PgdA. Clear conformational changes are predicted when the S protein is modeled in isolation or as part of a complex including PBP1a. (**B**) Host cells produce antimicrobials such as lysozyme and LL-37 causing bacterial cell envelope damage. (**C**) Paused diffusing and circumferentially moving PBP1a/PgdA/S protein complexes repair and modify PG after host-derived cell wall damage. At the site of damage, the S protein activates PBP1a and nascent PG is produced which is then the substrate of deacetylation by PgdA, rendering the newly repaired cell wall resistant to incoming host challenges. StkP and GpsB might also play a role in this regulation, as they interact with the intracellular moiety of S protein (**Fig. S4**). After repair, the complex is now competent to move to other sites requiring cell wall repair, while in absence of the S protein, the complex remains stuck.

Interestingly, PgdA and PBP1a of *L. monocytogenes*, a Gram-positive rod-shaped opportunistic human pathogen, distinct from streptococci, also interact, and their activities are controlled through GpsB^57^. We also identified a direct interaction between pneumococcal GpsB and the S protein, as well as with the serine-threonine kinase StkP (**Fig. S4**). Examining the *L. monocytogenes* genome for LysM-domain containing proteins, we identified Lmo1941, a protein of unknown function encoded near *cmk*, a conserved gene that flanks streptococcal *ess* genes (**Fig. S1**). Strikingly, a *lmo1941* knockout mutant was more susceptible to cephalosporins and showed an altered PG profile^58^. Similarly, the Gram-positive model organism *Bacillus subtilis* has a LysM domain-containing protein of unknown function, YpbE, which interacts with GpsB^55^. In addition, *B. subtilis* contains a PG deacetylase, PdaC, which confers reduced susceptibility to human lysozyme^59^. Future research should explore whether these S-like proteins also function together with PBP1a in non-streptococcal Gram-positive bacteria and investigate the hypothesis that the LysM domain of S proteins recognizes damaged PG, conveys this information to PBP1a, and what roles GpsB and StkP play in this regulation. It is noteworthy that the Alphafold prediction of the pneumococcal S protein shows large conformational changes when modeled in complex with PBP1a, suggesting that this flexibility might play a role in the mechanism for activating or coordinating PBP1a activity (**Fig. 7A**). The intrinsically disordered region (IDR) present within PBP1a might be important in this mechanism^60^.

A comprehensive understanding is emerging where streptococcal S proteins, and similar LysM domain-containing homologues in other Gram-positive bacteria, are key components of a complex involving a class A PBP, a PG deacetylase, and GpsB. This complex acts as a PG repair and modification system, functioning like sentinels scanning the cell wall for damage (**Fig. 7C**). In the absence of the S protein, this function is reduced or uncoordinated (**Fig. 5A**), leading to aberrant cell morphologies and increased susceptibility to cell wall damage (**Figs. 3, 6**). This repair/modification system is vital when bacteria encounter environmental challenges, such as cell wall-targeting antibiotics from competing microorganisms or antimicrobial peptides and enzymes like lysozyme from the host’s innate immune system (**Fig. 7B**). S proteins may thus represent as valuable therapeutic targets, such as for developing a new class of virulence inhibitors or as vaccine candidates^54^. It is interesting to note that the activities of major class A PBPs in Gram-negative bacteria are activated by the outer membrane lipoproteins LpoA and LpoB^61,62^. A functionally similar protein in *S. pneumoniae* is MacP, which promotes PG polymerization activity of PBP2a^63^. This raises the question of whether S proteins in Gram positive bacteria function analogously to MacP, activating PBP1a. Of note, simultaneous deletion of both *pbp1a* and *pbp2a* is lethal^64^ and in line with reduced PBP1a activity in a Δ*ess^Sp^* genetic background, we observed a significantly reduced growth rate in a Δ*pbp2a/*Δ*ess^Sp^* double mutant (Fig. S12). In addition, this hypothesis aligns with the observation that both the S protein and PBP1a are highly conserved in streptococci, whereas PgdA is less conserved (**Fig. 1, S1-2**), suggesting that the primary function of the S protein is linked to PBP1a.

## Methods

### Bacterial strains and culture conditions

All strains and the primers used are listed in **Table S6 & S7**. *S. pneumoniae* D39V strains were cultivated in liquid semi-defined C+Y medium, pH = 6.8^65^, at 37°C without shaking, from a starting optical density (OD_600nm_) of 0.01 until the desired OD. When not specified, the IPTG-inducible promoter (P_Lac_) was activated with 0.5 mM IPTG (Sigma Aldrich). When growing on solid media, Columbia agar (Sigma, 27688-500G) supplemented with 2.5% sheep blood at 37°C with 5% CO_2_ was used. When necessary, the medium was supplemented with the following antibiotics: erythromycin (ery) 0.5 μg.mL-1, kanamycin (kan) 250 μg.mL-1, spectinomycin (spc) 100 μg.mL-1, tetracycline (tet) 0.5 μg.mL-1, chloramphenicol (chl) 4 μg.mL−1, trimethoprim (trm) 8 μg.mL−1 and gentamycin (gen) 40 μg.mL−1. *S. pneumoniae* D39W strains were derived from unencapsulated strains IU1824 (D39W Δ*cps rpsL1*), which were derived from the encapsulated serotype-2 D39W progenitor strain IU1690^32,66^. Bacteria were cultured statically in Becton-Dickinson brain heart infusion (BHI) broth or C+Y at 37°C in an atmosphere of 5% CO_2_, and growth was monitored by OD_620_ as described^22^. For microscopy experiments, cells were inoculated from frozen glycerol stocks into BHI broth, serially diluted, and incubated 12–15 h statically at 37°C in an atmosphere of 5% CO_2_. The next day, 2-4 mL overnight cultures in BHI and still in exponential phase (OD_620_ = 0.1–0.4) were collected by centrifugation (21,100 x g for 5 min at room temperature (RT)), washed in 1 mL C+Y with centrifugation, resuspended in 4 mL fresh C+Y, and diluted to OD_620_ = 0.003-0.005 in 5 mL fresh C+Y. Cultures were sampled for phase contrast microscopy at OD_620_ ≈ 0.1-0.15. Unless noted, C+Y media were adjusted to pH = 7.1 by adding 500 µL of 1 M HCl to 43.9 mL of C+Y^65^. *Streptococcus salivarius* HSISS4 and its derivatives were grown at 37°C in M17G broth or CDMG without shaking^67^. *S. salivarius* transformation protocols was adapted from Mignolet 2018^67^. Briefly, cultures were started in the morning in CDMG and incubated for 3hr30min at 37°C. Then, the pheromone sComS (1µM final), as well as DNA (from overlapping PCRs), were added, and cells were allowed to recover for 3 hr at 37°C before plating on M17G agar supplemented with antibiotics when required. All synthetic media contained 1% glucose [w/v] (M17G and CDMG broth, respectively). When required, erythromycin (10 μg.mL-1) or xylose (1%) were added to the media. The synthetic peptide (purity of 95%) sComS, used for natural transformation (1 µM final concentration) was supplied by Peptide2.0 Inc. (Chantilly, VA, USA) and resuspended in DMSO.

### Strain construction

All *S. pneumoniae* D39V derivative strains were constructed by the integration of a linear DNA fragment containing up- and down-stream region of ∼1kb homology, into the chromosome by double homologous recombination. This linear DNA fragment was either obtained through a one-step PCR reaction of an existing strain containing the desired genomic modification or through Golden Gate cloning where different PCR products were ligated. Golden Gate cloning sites were introduced into the PCR fragments as part of the primers. PCR fragments were digested with either BsaI, Esp3I, SapI or AarI followed by ligation and transformation. Ligation product or genomic DNA were transformed into *S. pneumoniae* as previously described^65^, with cells taken at exponential growth phase (OD_600_ = 0.1) and induced with competence stimulation peptide 1 (CSP-1 for D39V and CSP-2 for TIGR4). Transformations were in all cases done in parental strains directly with the assembled products. Confirmation of the constructions were done by PCR and the resulting fragments were sequenced. All strains were stocked at OD_600_ ≈ 0.3 at -80°C with 14.5% of glycerol. The construction of all strains is detailed in **Table S6**. *S. pneumoniae* D39W derivative strains containing antibiotic markers were constructed by transformation of CSP1-induced competent pneumococcal cells with linear DNA amplicons synthesized by overlapping fusion PCR^22^. Primers and DNA templates used to synthesize different amplicons are listed in **Table S6 & S7**. Bacteria were grown on plates containing trypticase soy agar II (modified; Becton-Dickinson), and 5% (vol/vol) defibrinated sheep blood (TSAII-BA). TSAII-BA plates used for selection were incubated at 37°C in an atmosphere of 5% CO_2_, and contained antibiotics at concentrations described previously^21,22^. All constructs were confirmed by PCR and DNA sequencing of chromosomal regions corresponding to the amplicon region used for transformation. For *S. salivarius* transformation, *mNeonGreen* and *ess* genes were fused with overlapping PCRs to a strong (P_xyl1_) xylose-inducible promoter and inserted by double homologous recombination with the *xylR* regulator gene and an erythromycin resistance cassette at the permissive *gor* locus (downstream of HSISS4_00325).

### Conservation and gene neighborhood

S protein, PgdA, PBP1a and 16S sequences of different bacterial species were found using annotated genomes from ncbi.nlm.nih.gov. When possible, the sequence was confirmed using UniProt (www.uniprot.org). *S. pneumoniae* S protein sequence identity was then analyzed using p-BLAST (ncbi.nlm.nih.gov). 16S rRNA nucleotide sequences were used to generate a phylogenetic tree. Maximum likelihood method with a bootstrap of 1000 was performed using MEGA version 11^36^.

### Structure prediction using AlphaFold-Multimer

AlphaFold 3^68^ was used to generate the three-dimensional models of the proteins or complexes, using the AlphaFold Server (https://alphafoldserver.com/). ChimeraX was used to generate final models^69^.

### Phase contrast and fluorescence microscopy

Frozen stocks were inoculated 1:100 in C+Y medium (pH = 6.8) at 37°C until mid-exponential growth phase (OD_600_ = 0.2-0.3), supplemented with 100 µM IPTG when appropriate. Cultures were then diluted to OD_600_ = 0.01 and grow at 37°C until mid-exponential growth phase (OD_600_ = 0.2-0.3). Approx. 0.8 μL of culture were then used for imaging. The observation of morphological defect of cells exposed to beta-lactam antibiotic was done as followed: Frozen stocks were inoculated 1:100 in C+Y medium (pH = 6.8) at 37°C until mid-exponential growth phase (OD_600_ = 0.2-0.3). Cells were then diluted 100 fold and 0.006 µg.mL^-1^ of cefotaxime (CTX) was added to the cell culture when necessary. Cells were grown at 37°C for 1hr30min. They were then harvested by centrifugation (2 min at 9,000 x g), washed twice with 1 mL ice-cold PBS (pH = 7.4) and re-suspended in 25 uL ice-cold PBS. Cells were kept on ice prior to imaging. Cells were imaged by adding 0.8 µL of cell suspension onto 1.2% (w/v) PBS-agarose gel pads. Pads were then placed inside a gene frame (Thermo Fisher Scientific) and covered with a cover glass as described before^70^.

For preparation of agarose pads containing microholes with a diameter of 1.7 µm and height of 4 µm, we used a µVerCINI chip (Wunderlichips GmbH). For sample preparation, the mold was placed on top of 2% (w/v) melted PBS agarose until solidification. After removal of the mold, 10 µL of concentrated cell culture was loaded onto the microhole-agarose pad and centrifugation allowed the individual cells to enter the microholes. The pad was then covered with a cover glass.

Microscopy images were captured using a Leica DMi8 microscope with a DFC9000 GTC-VSC04862 camera, a HC PL APO 100x/1.40 oil objective and visualized using SOLA light engine (Lumencor®). Phase contrast images were acquired using transmission light (100 ms exposure). Filter sets used on the Leica DMi8 were the following: monomeric superfolder green fluorescent protein (msfGFP)/mNeonGreen (Ex: 470/40 nm Chroma ET470/40x, BS: LP 498 Leica 11536022, Em: 520/40 nm Chroma ET520/40m) with exposure time 700 ms, mScarlet-I-opt (Ex: Laser 550 nm Chroma ET545/30 X, BS: 595 nm Chroma 69008, Em: 635/70 nm Chroma ET635/70 m) with exposure time 700 ms. Images were acquired using LasX v.3.4.2.18368 (Leica).

Micrographs of *S. pneumoniae* D39W derivative strains were collected as follows: 500 µL of cultures in C+Y at OD_620_ ≈ 0.1-0.15 were centrifuged at 21,000 x g for 5 min at room temperature. 450 µL of supernatant was removed and the pellet was resuspended in the remaining 50 µL by gentle pipetting. 1 µL culture was placed on a glass slide, covered with a glass coverslip and imaged on a Nikon Eclipse E-400 microscope. Nikon NIS-Elements BR imaging software and a Nikon DS-Qi2 camera were used to capture images with a 20-50 ms exposure time.

*S. salivarius* cells were grown overnight in M17G medium at 37°C without any inducer. Cultures were then diluted to OD_600_ = 0.01 and grown with the addition of 1% xylose at 37°C until mid-exponential growth phase (OD_600_ = 0.3-0.4). 1 μL of culture was then used for imaging.

### Image analysis and cell segmentation

All microscopy images were analyzed using Fiji (fiji.sc). The deconvolution of the signals was performed, when appropriate, using Huygens (SVI) software with standard settings using 15 iterations. Cell segmentation based on phase-contrast images and fluorescent signals and demograph plots were performed using MicrobeJ^40^. Analysis and statistical comparison of cell size across different mutants and conditions was performed on R (version 4.3.0), using ggplot2 package. For every mutant and/or condition, cell size was recorded for at least 1,000 cells and their average cell length and width was determined. These averages were used to calculate the mean and SEM values that are shown in the main figures and the statistical differences were calculated using a Wilcoxon rank sum test. In Fig. 3G, we investigate the effect of antibiotic treatment on cell length in both WT strains (D39V vs D39V +CTX) and *ess* mutated strains (Δ*ess^Sp^*vs Δ*ess^Sp^* +CTX). In addition, we tested whether these differences are statistically significant between each other. Treated cells appear to be non-normally distributed. Therefore, we performed log_2_ base normalization followed by a two-way ANOVA. Lengths and widths of *Streptococcus pneumoniae* cells lacking capsule (**Fig. S6A**) were measured using Nikon NIS-Elements BR imaging software and compared using the nonparametric one-way ANOVA Kruskal-Wallis test with Dunn’s multiple comparisons test in GraphPad Prism.

### Time course experiments

Frozen stocks were inoculated 1:100 in C+Y medium (pH = 6.8) at 37°C until mid-exponential growth phase (OD_600_ = 0.1-0.2). Cultures were then diluted to OD_600_ = 0.003 and 250 µL of bacterial culture was then transferred in triplicate into 96-well plates. Growth was monitored by measuring the OD_600_ every 10 min for 20 hours at 37°C without CO_2_ in a TECAN Infinite F200 Pro. Approximately 0.8 μL of culture was taken at different time points for imaging directly in the 96-well plate. The experiment was done on two separate days.

### 3D Structured Illumination Microscopy (3D-SIM)

Samples for 3D-SIM were prepared as described above, by spotting 1 μL of culture onto PBS-10% acrylamide pads. DeltaVision OMX SR microscope (GE Healthcare) equipped with a 60x/1.42 NA objective (Olympus) and 488 nm and 568 nm excitation lasers, was used for the acquisition of the images. 16 Z-sections of 0.125 μm each were acquired in Structure Illumination mode with 20 ms exposure and 20% laser power. The images obtained were reconstructed with a Wiener constant of 0.001, and the volume reconstructed using SoftWoRx.

### Transmission Electron Microscopy (TEM)

For the TEM, frozen stocks of cells were inoculated 1:100 in 10 mL C+Y medium (pH = 6.8) at 37°C until OD_600_ = 0.2. Cells were then harvested by centrifugation (6 min at 4,000 x g) at 4°C and re-suspended in 2 mL of fresh C+Y medium. The culture was then transferred into a 2 mL Eppendorf tube and pelleted by centrifugation (1min30sec at 11,000 x g) at 4°C. The supernatant was washed out and cells were fixed in glutaraldehyde solution (EMS) 2.5% and in osmium tetroxide 1% (EMS) with 1.5% of potassium ferrocyanide (Sigma Aldrich) in phosphate buffer (PB 0.1 M [pH 7.4]) for 1 hr at room temperature. Cells were then washed twice with distilled water, then harvested by centrifugation (3,000 x g) and embedded in agarose (Sigma, St Louis, MO, US) 2% in water, dehydrated in acetone solution (Sigma Aldrich) at graded concentrations (30%-40 min; 70% - 40 min; 100% - 2x 1hr). This was followed by infiltration in Epon resin (EMS) at graded concentrations (Epon 33% in acetone-2 hr; Epon 66% in acetone-4 hr; Epon 100%-2x 8 hr) and finally polymerized for 48hr at 60°C in an oven. Ultra-thin sections of 50 nm were cut using a Leica Ultracut (Leica Mikrosysteme GmbH) and placed on a copper slot grid 2x 1mm (EMS) coated with a polystyrene film (Sigma Aldrich). Sections were post-stained with uranyl acetate (Sigma, St Louis, MO, US) 4% in water for 10 min, rinsed several times with water followed by Reynolds lead citrate in water (Sigma Aldrich) for 10 min and rinsed several times with water. Micrographs were taken with a transmission electron microscope JEOL JEM-2100Plus (JEOL Ltd.) at an acceleration voltage of 80kV with a TVIPS TemCam XF416 digital camera (TVIPS GmbH).

### Single-molecule dynamics

Single-molecule dynamics experiments were performed as described previously^24^, but are briefly described here.

#### Cell growth

Frozen glycerol stocks were inoculated into 4 mL brain heart infusion broth (BHI; BD Bacto 237500), serially diluted and incubated 12-16 hr at 37°C with 5% CO_2_ without shaking. 2-4 mL cultures with OD_620_ 0.1-0.4 were centrifuged (throughout procedure, all centrifuge steps were at 21,100×g for 5 min at room temperature) and washed in 1 mL C+Y (when testing strains containing a zinc-inducible promoter, all C+Y was supplemented with 0.25 mM ZnCl2 and 0.025 mM MnSO4 throughout the entire procedure). Samples were centrifuged again and resuspended in 4 mL fresh C+Y, then diluted to OD_620_ 0.003-0.005 in 5 mL fresh C+Y and incubated at 37°C with 5% CO_2_ without shaking.

#### HaloTag ligand labeling

At OD_620_ 0.08-0.15, 100-150 pM JF549 Halotag ligand (gift of Luke Lavis, Janelia) was added to 500 µL culture. Samples were incubated 15 min at 37°C with 400 rpm shaking in the dark, then centrifuged and washed in 0.9 mL C+Y. Centrifugation and washing was repeated 2 more times, then samples were centrifuged again and cell pellets were resuspended in 500 µL C+Y.

#### TIRFm sample preparation

Glass microscopy slides were cleaned with 70% ethanol and Gene Frames (1 cm x 1 cm; Thermo-Fisher, AB0576) were attached. 1.5% w/v agarose melted in C+Y was allowed to solidify inside the Gene Frames, creating a pad. 1.2 µL of samples were spread on a glass coverslip, briefly air dried (<3 m) and affixed with cells facing down onto the agarose pad. Slides were equilibrated at room temperature for 10 min, then incubated at 37°C for at least 20 min before imaging.

#### TIRFm imaging

TIRFm was performed in the Indiana University Light Microscopy Imaging Center on a DeltaVision OMX-SR (GE Healthcare) microscope with an Apo N 60X/1.49 TIRF objective (Olympus) and PCO.edge 4.2 sCMOS cameras (PCO). Differential Interference Contrast (DIC) images were used to determine cell locations. For HaloTag imaging, the laser line was 561 nm with emission filter 609-654. DIC light power was 10% (T), exposure time was 9 ms. HaloTag laser power was 100%, exposure time 45 msec. One image was acquired each second in sequential mode for a total of 180 sec, and channels were aligned using SoftWoRx (GE Healthcare).

#### TIRFm image analysis

Data were processed and analyzed using FIJI^71^. DIC signal was inverted to display dark cell bodies with light cell boundaries. If needed, the brightness of DIC frames were adjusted using the Enhance Contrast function. The plugin HyperstackReg^72^ was used to account for drift of cells during imaging, using rigid-body registration on the DIC channel. Movement pattern analysis determined the relative frequency of molecules moving diffusively, circumferentially, or static (non-moving). For each field, each cell was visualized throughout the full 180 sec of imaging, and HaloTag-PBP1a molecules were identified as having an approximate diameter of 3-6 pixels and a signal intensity greater than ≈ 1.2-fold above background. Only fields with >80% of cells having 0 or 1 HaloTag molecules were analyzed. Partially visible cells on the edge of a field were excluded from analysis. Circumferential molecules were defined as moving in one direction for ≥6 consecutive frames, with a linear velocity ≥5 nm/s as determined by kymograph analysis (see below). Static molecules were defined as motionless for ≥6 consecutive frames, or moving with a linear velocity <5 nm/s as determined by kymograph analysis. Diffusive molecules were defined as particles that moved, but not in a consistent direction for ≥6 frames (non-consecutive) within a period of 90 sec. A cell can have molecules displaying all 3 motion types, but no more than 1 molecule of each type can be scored per cell (for example, if 2 circumferential and 1 diffusive molecules are seen in a cell, only 1 circumferential and 1 diffusive molecule would be scored). Molecules were determined to be either within an area of the cell where PG synthesis occurs (midcell) or not (non-midcell). Before starting kymograph analysis, the spatial scale was removed from the images. The Line tool and Reslice function in FIJI were used to generate kymographs. Linear velocities were calculated using V = P / (T * tan(A)), where V is the linear velocity (in nm/sec), P is the pixel size (78.6 nm), T is the time between frames (in sec), tan is the tangent function, and A is the angle (in radians) of the molecule path measured from the kymograph. Molecule durations were measured by drawing a vertical line from the beginning to the end of the molecule path in the kymograph. For images acquired every second, the duration (in sec) equals the length of the line.

The relative frequencies of motion types were compared between strains using either an unpaired t test or a two-way ANOVA with a Newman-Keuls test (merodiploid strains) (GraphPad Prism). Velocities and durations were compared between strains using either a Kruskal-Wallis test with Dunn’s multiple comparisons test or Mann-Whitney test (merodiploid strains) (GraphPad Prism).

### Microtiter plate-based growth assay

*S. pneumoniae* growth assays were performed by first growing the frozen cells in C+Y medium (pH = 6.8) at 37°C until mid-exponential growth phase (OD_600_ = 0.2-0.3), supplemented with 100 µM IPTG when appropriate. Cultures were then diluted 100-fold into C+Y medium supplemented or not with IPTG and with or without the addition of antibacterial compounds (e.g. different concentrations of cell wall targeting antibiotics), as indicated in the main text and/or figures and/or figure legends. 250 µL of bacterial culture was then transferred in triplicate (technical replicate) into 96-well plates. Growth was monitored by measuring the OD_600_ every 10 min for 20 hr at 37°C without CO_2_ in a TECAN Infinite F200 Pro. OD_600_ values were normalized so that the lowest value measured during the first hour of growth was 0.004, the initial OD_600_ value of the inoculum. Each growth assay was performed in triplicate (biological replicate) and the mean value plotted, with the SEM (Standard Error of the Mean) as shading using BactExtract^73^.

### Proteomics analysis

All raw MS data together with raw output tables are available via the Proteomexchange data repository (www.proteomexchange.org) with the accession PXD055534.

#### Sample preparation and protein digestion

The GFP–S Protein and P3–GFP (negative control) strains were grown in C+Y medium at 37°C until OD_600_ = 0.2 and cells were harvested by centrifugation 15 min at 3,000 x g at 4°C. Cells were then incubated in lysis buffer (0.1 M Tris–HCl pH = 7.5, 150mM potassium acetate, 0.1% protease inhibitor cocktail (Sigma-Aldrich), 100 μg.mL−1 RNase A, 10 μg.mL−1 DNase (Sigma-Aldrich), 0.025% Deoxycholate, 5% glycerol and 1% n-Dodecyl-beta-Maltoside Detergent) for 20 min at 37°C. This was followed by 5 min sonication at 40% amplitude 1sec ON 2sec OFF with MS-73 microtip in ice. Cell debris were eliminated by centrifugation 30 min at 15,000 x g and 4°C. Cell lysate was then incubated with equilibrated GFP-Trap resin (Chromotek) at 4 °C for 2 hr. After several washes with wash buffer (10 mM Tris–HCl pH = 7.5, 150 mM NaCl, 0.5 mM EDTA), beads were stored at -80°C.

The samples were digested following a modified version of the iST method^74^ (named miST method). 25 µL of miST lysis buffer (1% Sodium deoxycholate, 100mM Tris pH = 8.6, 10 mM DTT), were added to the beads. After mixing and dilution 1:1 (v:v) with H2O, samples were heated 5 min at 75°C. Reduced disulfides were alkylated by adding 13 µL of 160 mM chloroacetamide (33 mM final) and incubating for 45 min at 25°C in the dark. After digestion with 1.0 μg of Trypsin/LysC mix (Promega #V5073) for 2 hr at 25°C, sample supernatants were transferred in new tubes. To remove sodium deoxycholate, two sample volumes of isopropanol containing 1% TFA were added to the digests, and the samples were desalted on a strong cation exchange (SCX) plate (Oasis MCX; Waters Corp., Milford, MA) by centrifugation. After washing with isopropanol/1%TFA, peptides were eluted in 200 µL of 80% MeCN, 19% water, 1% (v/v) ammonia, dried by centrifugal evaporation and redissolved in 2% acetonitrile, 0.1% formic acid.

#### Liquid Chromatography-Mass Spectrometry analyses

##### Exploris 480

Tryptic peptide mixtures were injected on a Ultimate RSLC 3000 nanoHPLC system interfaced via a nanospray Flex source to a high resolution Orbitrap Exploris 480 mass spectrometer (Thermo Fisher, Bremen, Germany). Peptides were loaded onto a trapping microcolumn Acclaim PepMap100 C18 (20 mm x 100 μm ID, 5 μm, Dionex) before separation on a C18 custom packed column (75 μm ID × 45 cm, 1.8 μm particles, Reprosil Pur, Dr. Maisch), using a gradient from 4 to 90% acetonitrile in 0.1% formic acid for peptide separation (total time: 140 min). Full MS survey scans were performed at 120,000 resolution. A data-dependent acquisition method controlled by Xcalibur software (Thermo Fisher Scientific) was used that optimized the number of precursors selected (“top speed”) of charge 2+ to 5+ while maintaining a fixed scan cycle of 2 s. Peptides were fragmented by higher energy collision dissociation (HCD) with a normalized energy of 30% at 15’000 resolution. The window for precursor isolation was of 1.6 m/z units around the precursor and selected fragments were excluded for 60s from further analysis.

#### Data processing

##### Mascot (DDA)

MS data were analyzed using Mascot 2.7 (Matrix Science, London, UK) set up to search the *S. pneumoniae* reference proteome (RefProt) based on the UniProt database (www.uniprot.org), version of May 3rd, 2020, containing 1’915 sequences), and a custom contaminant database containing the most usual environmental contaminants and enzymes used for digestion (keratins, trypsin, etc.). Trypsin (cleavage at K,R) was used as the enzyme definition, allowing 2 missed cleavages. Mascot was searched with a parent ion tolerance of 10 ppm and a fragment ion mass tolerance of 0.02 Da. Carbamidomethylation of cysteine was specified in Mascot as a fixed modification. Protein N-terminal acetylation and methionine oxidation were specified as variable modifications.

#### Data analysis

##### Scaffold (DDA)

Scaffold (version Scaffold 5.2.2, Proteome Software Inc., Portland, OR) was used to validate MS/MS based peptide and protein identifications. Peptide identifications were accepted if they could be established at greater than 90.0% probability by the Percolator posterior error probability calculation^75^. Protein identifications were accepted if they could be established at greater than 95.0% probability and contained at least 2 identified peptides. Protein probabilities were assigned by the Protein Prophet algorithm^76^. Proteins that contained similar peptides and could not be differentiated based on MS/MS analysis alone were grouped to satisfy the principles of parsimony. Proteins sharing significant peptide evidence were grouped into clusters.

### Split-luciferase assay

Frozen *S. pneumoniae* cells were grown in C+Y medium with 0.1mM IPTG when needed at 37 °C until OD_600_ = 0.3. 1% NanoGlo Live Cell substrate (Promega) was then added and 300 µL of bacterial culture was then transferred in triplicate into 96-well plates. Luminescence was measured then 15 times at 37 °C every 30 sec in a plate reader (TECAN Infinite F200 Pro). For each technical replicate, we selected the minimal value of n = 15 measurements. The relative luminescent unit (RLU) value represents the log10+1 of the average of the technical (n = 3) and biological (n≥ 3) replicates. Finally, we normalized the values by subtracting D39V value to all other values. Minus values were set to zero. Heatmaps were generated using R (version 4.3.0) and the ggplot2 package

### CRISPRi-seq screening

#### Construction of the knockout background strains

The S protein deletion (*ess*::*kan*) was amplified from the clean deletion strains previously made and transformed into *S. pneumoniae* D39V strain DCI23^77^, allowing the CRISPRi system to operate. Constructs were checked by PCR and sequencing.

#### Insertion of the pooled CRISPRi library in *S. pneumoniae* D39V

To construct an IPTG-inducible CRISPRi-seq libraries, 1 ng.µL-1 of a pooled sgRNA library targeting 1499 pneumococcus operons was transformed into VL3744 and VL1998 (**Table S6**)^77^. The transformants were plated using beads on three 25 mL Columbia blood agar plates with 100 µg. mL-1 of spectinomycin, to have enough coverage. After 17 hr of growth at 37°C with 5% CO_2_, two plates of each transformant were harvested by washing with fresh C+Y medium containing 100 µg.mL-1 of spectinomycin and stocked at -80°C with 20% glycerol. From this stock, a preculture was made, where 400 µL of cells were inoculated in 4.5 mL of C+Y medium containing 100 µg.mL-1 of spectinomycin, up to OD_600_ = 0.3, and stocked at -80°C with 20% glycerol.

#### CRISPRi-seq screen induction

About 21 generations of growth, done in quadruplicate, were performed where the CRISPRi system was induced and cells were in competition. To do so, 50 µL of each preculture was inoculated in 2 mL of C+Y medium containing 100 µg.mL-1 of spectinomycin and IPTG 100 µM. The same was done without inducing the CRISPRi system, thus without IPTG. When the OD_600_ = 0.25, 50 µL cultures were diluted into 10 mL of fresh medium. Bacteria were then collected when the OD_600_ = 0.4, and the pellet of each tube were used for genomic DNA extraction. gDNA concentration as measured using NanoDrop 2000c (ThermoFisher).

#### Library preparation for Illumina MiniSeq sequencing

Illumina libraries were prepared using a one-step PCR, with the isolated gDNA as template. 2 µL of primers and 4 µg of gDNA were used for each 40 µL PCR reaction, all prepared on ice. The PCR elongation time was set at 72°C for 1 min and 8 cycles were performed. The PCR amplicons (303bp) were then purified from a 2% agarose gel, and the amount of DNA was quantified by Qubit assay (Q33227, ThermoFisher Scientific). The purified libraries were then pooled together following Illumina protocol (“Denature and Dilute Libraries Guide”, document #1000000002697 v02), and sequenced using an Illumina MiniSeq High Output Reagent Kit. To prevent cluster calling, the first 54 of the 150 cycles of the sequencing were not considered.

#### Data analysis

Using 2FAST2Q^78^, sgRNA read count was calculated for each condition, by mapping the reads to all 1499 sgRNA sequences. More precisely, for each read, everything except the 20 base-pairing regions of sgRNA were trimmed off, PhreD of 30 and one mismatch allowed max were set as parameters. The output of the program was then analyzed using R (version 4.3.0), with an adapted version of a previous script (https://github.com/veeninglab/CRISPRi-seq^44^), which used the DESeq2 package for evaluation of the fitness cost of each sgRNA. The p-values were adjusted using False Discovery Rate (FDR) and the threshold was set at 0.1. We tested against a log2FC of 1. Plots where generated using R.

### Genome resequencing of multiple mutants by NGS

Strains mentioned in **Table S6** were grown in C+Y medium at 37°C until OD_600_ = 0.3 and then harvested by centrifugation 1 min at 10,000 x g for genomic DNA isolation using FastPure Bacteria, DNA isolation Mini Kit (Vazyme). The extracted genomes were then analyzed by Illumina sequencing PE150 (Microbial whole genome library preparation (350bp) (Novogene). Raw reads were ranging between 7,761,672 and 18,822,346. Mutations were mapped onto D39V genome using breseq pipeline^79^.

### Persister phenotype of *S. pneumoniae*

Streptococcal isolates were thawed from -80°C 17% glycerol stocks, inoculated and pre-cultured in 2 mL of sterile nutrient rich growth medium (45% M17, 45% CAT [i.e. 10 g casamino acids (Difco, cat no. 228830), 5 g tryptone (Oxoid, cat no. LP0042B), 5g sodium chloride, and 10 g yeast extract (Oxoid, cat no. LP0021B) per L sterile water], 0.225% glucose, and 26 U/mL catalase (Sigma-Aldrich, cat no. C1345) in 0.05M K2HPO4) at 37°C to an OD_600_ = 0.3. In a sterile flat-bottomed 96-well plate (Nunclon Surface, Nunc, Denmark), 5 μL of pre-culture was inoculated to 195 μL of prewarmed growth medium. Bacterial growth at 37°C was monitored by measuring OD_595_ every 10 min in microplate reader without CO_2_ (Spark 10M, Tecan, Switzerland). When mid-exponential growth (OD_595_ = 0.15-0.25 for *S. pneumoniae)* was reached, penicillin G was added in a volume of 10 μL to an end concentration of 4 μg.mL-1. For penicillin-susceptible *S. pneumoniae* isolates, the minimum inhibitory concentrations (MICs) of penicillin G are ≤ 0.06 μg.mL^-1^ ^80^. Therefore, for susceptible isolates the applied concentration is ≥ 100x the MIC. Incubation at 37°C was immediately continued and OD_595_ monitored. Penicillin treatment induces autolysis and once OD_595_ had returned to blank medium baseline levels, 100 μL of fresh pre-warmed rich medium was added plus penicillinase (from *B. cereus*, Sigma-Aldrich, cat no. P0389) in 10 μL of 0.1M TRIS HCl pH = 7 + 0.1% BSA to an end concentration of 0.0125 units.mL^-1^. Incubation at 37°C was immediately continued and OD_595_ monitored every 10 minutes for another 24 hours. Afterwards, penicillin susceptibility was determined by spreading 10 μL from a well to a Columbia 5% blood agar plate using a sterile loop, placing a 1 μg oxacillin disk (BD, cat no. 254822) at the center, and measuring the zone of inhibition diameter after overnight culture at 37°C and 5% CO_2_. Growth curves were plotted using BactExtract^73^.

### PG analysis

#### Isolation of pneumococcal cell wall

Cell wall from *S. pneumoniae* was purified as described previously^49^ with modifications. Cells were grown in 1 L C+Y media at 37°C with 5% CO_2_ without shaking until the OD_600_ reached approximately 0.45. Cultures were harvested by centrifugation for 30 min at 4°C and 5,000 × *g*. The cell pellet was resuspended in 40 mL of ice-cold 50 mM Tris–HCl pH = 7.0. The cell suspension was added dropwise into a flask containing 120 mL of boiling 5% sodium dodecyl sulfate (SDS) solution with continuous stirring. The solution was boiled for another 15 min before cooling down to RT. The cell suspension was pelleted by ultracentrifugation at 25°C for 30 min at 130,000 × g. The pellet was washed twice with 20 mL of 1 M NaCl and repeatedly with deionized water (dH2O) until it was free of SDS, confirmed by a previously published assay^81^. The pellet was resuspended in 2 to 4 mL of dH2O, 1/3 volume of acid-washed glass beads (diameter of 0.17–0.18 mm, Sigma, München, Germany) was added, and cells were disrupted in a FastPrep machine (FP120, Thermo Scientific, Hemel Hempstead, UK). For cell disruption, 6 times 3 pulses at maximal speed were applied for 30 sec, with 5-min cooling periods on ice after 3 pulses. The glass beads were then separated from the cell lysate with a grade 4 glass filter (Winzer, Wertheim, Germany). The filtrate was centrifuged at 2000 × g for 5 min, and the supernatant, which contains the disrupted cell wall, was centrifuged at 130,000 × g for 30 min at 25°C. The pellet was resuspended in 10 mL of 100 mM Tris–HCl pH = 7.5 containing 20 mM MgSO_4_. DNase I and RNase I were added to final concentrations of 10 and 50 µg.mL^-1^, respectively, and the sample was stirred for 2 hr at 37°C. CaCl_2_ (10 mM) and trypsin (100 µg.mL^-1^) were added, and the sample was stirred for 18 hr at 37°C. Then, 1% SDS (final concentration) was added to have a final volume of 20-30 mL, and the sample was incubated for 15 min at 80°C to inactivate the enzymes. The cell wall was recovered by centrifugation for 30 min at 130,000 × g at 25°C, resuspended with 10 mL of 8 M LiCl, and incubated for 15 min at 37°C. After another centrifugation (see above), the pellet was resuspended in 10 mL of 100 mM ethylenediaminetetraacetic acid (EDTA) pH = 7.0 and incubated at 37°C for 15 min. The cell wall was washed twice with 20-30 dH2O. The pellet was resuspended pellet in 2 to 4 mL of dH2O and lyophilized. Cell wall samples (typically 12-15 mg from 1-L culture) were stored at - 20°C.

#### Preparation of peptidoglycan

Five mg of cell wall was stirred with 3 mL of 48% hydrofluoric acid (HF) for 48 h at 4°C in an ultracentrifuge tube made of polyallomer tightly closed by Parafilm. The volatile, aggressive, and toxic HF was handled with great caution to avoid any contact with the skin and spillage. The peptidoglycan was recovered by centrifugation (45 min, 130,000 × g, 4°C) and washed with ice-cold dH2O, 100 mM Tris–HCl pH = 7.0, and then twice with dH2O. The pellet containing 2 to 2.5 mg of peptidoglycan was resuspended in 500 µl of 0.02% NaN3 and stored at 4°C.

#### Preparation of muropeptides

To release the muropeptides from peptidoglycan, 180 µL of murein preparation (above) was mixed with 60 µL of 4× digestion buffer (80 mM sodium phosphate pH = 4.8). Then, 40 µg.mL^-1^ of cellosyl was added and the mixture was stirred overnight at 37°C. The addition of cellosyl and incubation was repeated. The reaction was stopped by heating the sample for 10 min at 100°C. The sample was cleared by centrifugation for 10 min at 13,000 × g. For reduction, 1 sample volume of 500 mM sodium borate pH 9.0 and a few crystals of NaBH_4_ were added, followed by 30 min of incubation at 20°C. The reaction was stopped by adjusting the pH to 3.5 to 4.5 with 20% H_3_PO_4_.

#### Separation of muropeptides by HPLC

Muropeptides were separated on a reversed-phase column (Prontosil 120-3-C18-AQ, 250 × 4.6 mm, 3 µM, Bischoff, Germany) using an Agilent 1100 system operating ChemStation software. HPLC was performed with a column temperature of 55°C using a linear 135-min gradient from 100% buffer A (10 mM sodium phosphate pH 6.0 and 20 µM NaN_3_) to 100% buffer B (10 mM sodium phosphate pH 6.0 and 30% methanol) (Fisher Scientific, Loμghboroμgh, UK). Muropeptide fractions were detected at 205 nm. The muropeptides were quantified by integration of their peak area using Laura 4 software (LabLogic, Sheffield, UK). The total peak area from all fractions, excluding the salt fractions (eluting before 8 min), was normalized to 100%, and the relative area of each fraction was determined.

### Lysozyme and LL-37 susceptibility assay

To test the sensibility of *S. pneumoniae* strains to lysozyme and LL-37, frozen cells were first grown in C+Y medium (pH = 6.8) at 37°C until mid-exponential growth phase (OD_600_ = 0.2-0.3). Strains were then diluted 10 fold into C+Y medium in different concentrations of lysozyme (0, 20, 40 or 80 μg.mL-1) and LL-37 (0, 10, 20 or 30 μg.mL-1). 250 µL of bacterial culture was then added in triplicate in a 96-well microtiter plate reader where the cellular growth (OD_600_) was monitored every 10 min for 20 hours at 37°C without CO_2_ in a TECAN Infinite F200 Pro. The experiment was done three times. Here, the heatmaps of the area under the curve (AUC) represent one of the biological triplicates. We calculated the empiric AUC based on the optical density (OD_500_), by summing together the first 7 hours of growth (which correspond to the time needed for the slowest strain to reach its highest OD_500_) using BactExtract^73^.

### Zebrafish husbandry and maintenance

Transparent adult casper mutant zebrafish (*mitfa^w2/w2^*;*roy^a9/a9^*) were maintained at 26°C in aerated 5-L tanks with a 10/14 hr dark/light cycle^82^. Zebrafish embryos were collected within the first hours post fertilization (hpf) and kept at 28°C in E3 medium (5.0 mM NaCl, 0.17 mM KCl, 0.33 mM CaCl·2H_2_O, 0.33 mM MgCl_2_·7H_2_O) supplemented with 0.3 mg.L-1 methylene blue. All procedures related to zebrafish studies at UNIL comply with the Swiss regulations on animal experimentation (Animal Welfare Act SR 455 and Animal Welfare Ordinance SR 455.1).

### Zebrafish survival experiments

*S*. *pneumoniae* D39V wild type strain or S protein mutated strain (D39V, Δ*ess^Sp^*) were grown in C+Y medium at 37°C until mid-exponential growth phase (OD_600_ = 0.2). Cells were harvested by centrifugation 10 min at 6,000 x g, the supernatant removed and resuspended in 0.125% (w/v) phenol red solution (Sigma-Aldrich; P0290) to aid visualization of the injection process. Prior to injection, 2 days post fertilization (dpf) larvae were mechanically dechorionated if necessary and anesthetized in 0.02% (w/v) Tricaine (Sigma-Aldrich, Cat# A5040). Larvae were randomly assigned to experimental groups. The bacterial suspension was then injected into the hindbrain ventricle of *casper* zebrafish larvae at 2 dpf. After injection, the larvae were kept in 6-well plates containing egg water (60 μg.mL-1 sea salts (Sigma-Aldrich; S9883) in demi water) at 28°C and the mortality rate monitored at fixed time-points as described previously^83^. All experiments were performed in triplicate. Survival graphs were generated with GraphPad Prism 10.2.1 and analyzed with the log rank (Mantel-Cox) test. Results were considered significant at p values < 0.05.

### Mouse experiments

All animal studies were conducted in accordance with the Animal Welfare Act and approved protocols recommended by the University of California San Diego Institutional Animal Care and Use Committee (IACUC, #S00227M). Mice were housed in a 12 hr light/dark cycle in a specific pathogen free facility. Mice were housed in prebedded corn cob disposable cages and received acidified water and 2020X diet. 8-12 week old male and female C57BL/6 (Jackson Labs, 000664) were infected intratracheally with ∼5*10^6^ CFUs D39V, Δ*ess^Sp^* mutant or P*_lac_-ess* Δ*ess^Sp^* complemented *S. pneumoniae* strains. Mice were anesthetized with 100 mg.kg-1 ketamine and 10 mg.kg-1 xylazine intraperitoneally before injection and monitored on a heating pad until fully recovered from anesthesia. Mice were monitored twice daily for morbidity and mortality for seven days. Survival graphs were generated with GraphPad Prism 10.2.1 and analyzed with the log rank (Mantel-Cox) test.

## Data availability

The data that support the findings of this study are incorporated in the manuscript and its supporting information. Genome sequences, assemblies, and sequencing reads are available at NCBI (BioProject accession number pending).

## Declaration of interests

The authors declare no competing interests

## Acknowledgements

We thank the members of the Veening group for valuable discussions, Florian Patrick Bock for help with protein structure prediction and Vincent de Bakker for help with CRISPRi-seq data analysis. We thank the UNIL GTF for sequencing, PAF for proteomics and EMF for TEM. We thank Nadine Vastenhouw for access to the zebrafish facility and continued support. We thank Dirk van Swaay (Wünderlichips) for design and production of the micropillars chip. We thank David Gonzalez for strains and insightful discussions. Work in the lab of J.W.V. was supported by SNSF grants 310030_192517, 310030_200792 and NCCR 51NF40_180541. Work in the lab of M.E.W. was supported by NIH grant R35GM131767 and NIH equipment grant S10OD024988 to the Indiana University Bloomington Light Microscopy Imaging Center. W.V. was supported by the UK Biotechnology and Biological Sciences Research Council (BBSRC; BB/W013630/1).

## Author contributions

J.B. and J.W.V. wrote the paper with input from all authors. J.B., C.G., K.B., E.B, L.M., K.J., H.-C.T.T., A.K., J.M., D.V., I.W. performed the experiments. J.B., C.G., K.B., E.B., K.J., H.-C.T.T., A.C., J.M., D.V., J.B., I.W., V.N., D.G., W.V., M.E.W. and J.-W.V. designed, analyzed and interpreted the data.

## Competing interests

The authors declare no competing interests.

## Notes

### Competing Interest Statement

The authors have declared no competing interest.

## References

1. Abranches, J., Zeng, L., Kajfasz, J.K., Palmer, S.R., Chakraborty, B., Wen, Z.T., Richards, V.P., Brady, L.J., and Lemos, J.A. (2018). Biology of Oral Streptococci. Microbiol. Spectr. 6. 10.1128/microbiolspec.GPP3-0042-2018.

2. Martinović, A., Cocuzzi, R., Arioli, S., and Mora, D. (2020). *Streptococcus thermophilus*: To Survive, or Not to Survive the Gastrointestinal Tract, That Is the Question! Nutrients 12, 2175. 10.3390/nu12082175.

3. Kampff, Z., van Sinderen, D., and Mahony, J. (2023). Cell wall polysaccharides of streptococci: A genetic and structural perspective. Biotechnol. Adv. 69, 108279. 10.1016/j.biotechadv.2023.108279.

4. Carapetis, J.R., Steer, A.C., Mulholland, E.K., and Weber, M. (2005). The global burden of group A streptococcal diseases. Lancet Infect. Dis. 5, 685–694. 10.1016/S1473-3099(05)70267-X.

5. Ralph, A.P., and Carapetis, J.R. (2013). Group a streptococcal diseases and their global burden. Curr. Top. Microbiol. Immunol. 368, 1–27. 10.1007/82_2012_280.

6. Sims Sanyahumbi, A., Colquhoun, S., Wyber, R., and Carapetis, J.R. (2016). Global Disease Burden of Group A Streptococcus. In: Streptococcus pyogenes: Basic Biology to Clinical Manifestations.

7. Guerra, S., and LaRock, C. (2024). Group A Streptococcus interactions with the host across time and space. Curr. Opin. Microbiol. 77, 102420. 10.1016/j.mib.2023.102420.

8. Ling, J., and Hryckowian, A.J. (2024). Re-framing the importance of Group B Streptococcus as a gut-resident pathobiont. Infect. Immun., e0047823. 10.1128/iai.00478-23.

9. GBD 2021 Lower Respiratory Infections and Antimicrobial Resistance Collaborators (2024). Global, regional, and national incidence and mortality burden of non-COVID-19 lower respiratory infections and aetiologies, 1990-2021: a systematic analysis from the Global Burden of Disease Study 2021. Lancet Infect. Dis., S1473-3099(24)00176–2. 10.1016/S1473-3099(24)00176-2.

10. Mistou, M.-Y., Sutcliffe, I.C., and van Sorge, N.M. (2016). Bacterial glycobiology: rhamnose-containing cell wall polysaccharides in Gram-positive bacteria. FEMS Microbiol. Rev. 40, 464–479. 10.1093/femsre/fuw006.

11. Vollmer, W., Massidda, O., and Tomasz, A. (2019). The Cell Wall of *Streptococcus pneumoniae*. Microbiol. Spectr. 7. 10.1128/microbiolspec.GPP3-0018-2018.

12. Weiser, J.N., Ferreira, D.M., and Paton, J.C. (2018). *Streptococcus pneumoniae*: transmission, colonization and invasion. Nat. Rev. Microbiol. 16, 355–367. 10.1038/s41579-018-0001-8.

13. Gibson, P.S., and Veening, J.-W. (2023). Gaps in the wall: understanding cell wall biology to tackle amoxicillin resistance in *Streptococcus pneumoniae*. Curr. Opin. Microbiol. 72, 102261. 10.1016/j.mib.2022.102261.

14. Mills, J.O., and Ghosh, P. (2021). Nonimmune antibody interactions of Group A Streptococcus M and M-like proteins. PLoS Pathog. 17, e1009248. 10.1371/journal.ppat.1009248.

15. Wierzbicki, I.H., Campeau, A., Dehaini, D., Holay, M., Wei, X., Greene, T., Ying, M., Sands, J.S., Lamsa, A., Zuniga, E., et al. (2019). Group A Streptococcal S Protein Utilizes Red Blood Cells as Immune Camouflage and Is a Critical Determinant for Immune Evasion. Cell Rep. 29, 2979–2989.e15. 10.1016/j.celrep.2019.11.001.

16. Campeau, A., Uchiyama, S., Sanchez, C., Sauceda, C., Nizet, V., and Gonzalez, D.J. (2021). The S Protein of Group B Streptococcus Is a Critical Virulence Determinant That Impacts the Cell Surface Virulome. Front. Microbiol. 12, 729308. 10.3389/fmicb.2021.729308.

17. Straume, D., Piechowiak, K.W., Kjos, M., and Håvarstein, L.S. (2021). Class A PBPs: It is time to rethink traditional paradigms. Mol. Microbiol. 116, 41–52. 10.1111/mmi.14714.

18. Land, A.D., Tsui, H.-C.T., Kocaoglu, O., Vella, S.A., Shaw, S.L., Keen, S.K., Sham, L.-T., Carlson, E.E., and Winkler, M.E. (2013). Requirement of essential Pbp2x and GpsB for septal ring closure in *Streptococcus pneumoniae* D39. Mol. Microbiol. 90, 939–955. 10.1111/mmi.12408.

19. Massidda, O., Nováková, L., and Vollmer, W. (2013). From models to pathogens: how much have we learned about *Streptococcus pneumoniae* cell division? Environ. Microbiol. 15, 3133–3157. 10.1111/1462-2920.12189.

20. Philippe, J., Vernet, T., and Zapun, A. (2014). The elongation of ovococci. Microb. Drug Resist. Larchmt. N 20, 215–221. 10.1089/mdr.2014.0032.

21. Tsui, H.-C.T., Boersma, M.J., Vella, S.A., Kocaoglu, O., Kuru, E., Peceny, J.K., Carlson, E.E., VanNieuwenhze, M.S., Brun, Y.V., Shaw, S.L., et al. (2014). Pbp2x localizes separately from Pbp2b and other peptidoglycan synthesis proteins during later stages of cell division of *Streptococcus pneumoniae* D39. Mol. Microbiol. 94, 21–40. 10.1111/mmi.12745.

22. Tsui, H.-C.T., Zheng, J.J., Magallon, A.N., Ryan, J.D., Yunck, R., Rued, B.E., Bernhardt, T.G., and Winkler, M.E. (2016). Suppression of a deletion mutation in the gene encoding essential PBP2b reveals a new lytic transglycosylase involved in peripheral peptidoglycan synthesis in *Streptococcus pneumoniae* D39. Mol. Microbiol. 100, 1039–1065. 10.1111/mmi.13366.

23. Lamanna, M.M., Manzoor, I., Joseph, M., Ye, Z.A., Benedet, M., Zanardi, A., Ren, Z., Wang, X., Massidda, O., Tsui, H.-C.T., et al. (2022). Roles of RodZ and class A PBP1b in the assembly and regulation of the peripheral peptidoglycan elongasome in ovoid-shaped cells of *Streptococcus pneumoniae* D39. Mol. Microbiol. 118, 336–368. 10.1111/mmi.14969.

24. Perez, A.J., Lamanna, M.M., Bruce, K.E., Touraev, M.A., Page, J.E., Shaw, S.L., Tsui, H.-C.T., and Winkler, M.E. (2024). Elongasome core proteins and class A PBP1a display zonal, processive movement at the midcell of *Streptococcus pneumoniae*. Proc. Natl. Acad. Sci. U. S. A. 121, e2401831121. 10.1073/pnas.2401831121.

25. Straume, D., Piechowiak, K.W., Olsen, S., Stamsås, G.A., Berg, K.H., Kjos, M., Heggenhougen, M.V., Alcorlo, M., Hermoso, J.A., and Håvarstein, L.S. (2020). Class A PBPs have a distinct and unique role in the construction of the pneumococcal cell wall. Proc. Natl. Acad. Sci. U. S. A. 117, 6129–6138. 10.1073/pnas.1917820117.

26. Morè, N., Martorana, A.M., Biboy, J., Otten, C., Winkle, M., Serrano, C.K.G., Montón Silva, A., Atkinson, L., Yau, H., Breukink, E., et al. (2019). Peptidoglycan Remodeling Enables *Escherichia coli* To Survive Severe Outer Membrane Assembly Defect. mBio 10, e02729–18. 10.1128/mBio.02729-18.

27. Vigouroux, A., Cordier, B., Aristov, A., Alvarez, L., Özbaykal, G., Chaze, T., Oldewurtel, E.R., Matondo, M., Cava, F., Bikard, D., et al. (2020). Class-A penicillin binding proteins do not contribute to cell shape but repair cell-wall defects. eLife 9, e51998. 10.7554/eLife.51998.

28. Vollmer, W., and Tomasz, A. (2000). The pgdA gene encodes for a peptidoglycan N-acetylglucosamine deacetylase in *Streptococcus pneumoniae*. J. Biol. Chem. 275, 20496–20501. 10.1074/jbc.M910189199.

29. Cole, A.M., Liao, H.-I., Stuchlik, O., Tilan, J., Pohl, J., and Ganz, T. (2002). Cationic polypeptides are required for antibacterial activity of human airway fluid. J. Immunol. Baltim. Md 1950 169, 6985–6991. 10.4049/jimmunol.169.12.6985.

30. Vollmer, W., and Tomasz, A. (2002). Peptidoglycan N-acetylglucosamine deacetylase, a putative virulence factor in *Streptococcus pneumoniae*. Infect. Immun. 70, 7176–7178. 10.1128/IAI.70.12.7176-7178.2002.

31. Davis, K.M., Akinbi, H.T., Standish, A.J., and Weiser, J.N. (2008). Resistance to mucosal lysozyme compensates for the fitness deficit of peptidoglycan modifications by *Streptococcus pneumoniae*. PLoS Pathog. 4, e1000241. 10.1371/journal.ppat.1000241.

32. Slager, J., Aprianto, R., and Veening, J.-W. (2018). Deep genome annotation of the opportunistic human pathogen *Streptococcus pneumoniae* D39. Nucleic Acids Res. 46, 9971–9989. 10.1093/nar/gky725.

33. Jumper, J., Evans, R., Pritzel, A., Green, T., Figurnov, M., Ronneberger, O., Tunyasuvunakool, K., Bates, R., Žídek, A., Potapenko, A., et al. (2021). Highly accurate protein structure prediction with AlphaFold. Nature 596, 583–589. 10.1038/s41586-021-03819-2.

34. Buist, G., Steen, A., Kok, J., and Kuipers, O.P. (2008). LysM, a widely distributed protein motif for binding to (peptido)glycans. Mol. Microbiol. 68, 838–847. 10.1111/j.1365-2958.2008.06211.x.

35. Whitley, K.D., Middlemiss, S., Jukes, C., Dekker, C., and Holden, S. (2022). High-resolution imaging of bacterial spatial organization with vertical cell imaging by nanostructured immobilization (VerCINI). Nat. Protoc. 17, 847–869. 10.1038/s41596-021-00668-1.

36. Tamura, K, Stecher, G., and Kumar, S. (2021). MEGA11: Molecular Evolutionary Genetics Analysis Version 11. Mol. Biol. Evol. 38. 10.1093/molbev/msab120.

37. Zuber, B., Haenni, M., Ribeiro, T., Minnig, K., Lopes, F., Moreillon, P., and Dubochet, J. (2006). Granular layer in the periplasmic space of gram-positive bacteria and fine structures of *Enterococcus gallinarum* and *Streptococcus gordonii* septa revealed by cryo-electron microscopy of vitreous sections. J. Bacteriol. 188, 6652–6660. 10.1128/JB.00391-06.

38. Oliveira Paiva, A.M., Friggen, A.H., Qin, L., Douwes, R., Dame, R.T., and Smits, W.K. (2019). The Bacterial Chromatin Protein HupA Can Remodel DNA and Associates with the Nucleoid in *Clostridium difficile*. J. Mol. Biol. 431, 653–672. 10.1016/j.jmb.2019.01.001.

39. Gallay, C., Sanselicio, S., Anderson, M.E., Soh, Y.M., Liu, X., Stamsås, G.A., Pelliciari, S., van Raaphorst, R., Dénéréaz, J., Kjos, M., et al. (2021). CcrZ is a pneumococcal spatiotemporal cell cycle regulator that interacts with FtsZ and controls DNA replication by modulating the activity of DnaA. Nat. Microbiol. 6, 1175–1187. 10.1038/s41564-021-00949-1.

40. Ducret, A., Quardokus, E.M., and Brun, Y.V. (2016). MicrobeJ, a tool for high throughput bacterial cell detection and quantitative analysis. Nat. Microbiol. 1, 16077. 10.1038/nmicrobiol.2016.77.

41. Sanchez-Puelles, J.M., Ronda, C., Garcia, J.L., Garcia, P., Lopez, R., and Garcia, E. (1986). Searching for autolysin functions. Characterization of a pneumococcal mutant deleted in the *lytA* gene. Eur. J. Biochem. 158, 289–293. 10.1111/j.1432-1033.1986.tb09749.x.

42. Beilharz, K., Nováková, L., Fadda, D., Branny, P., Massidda, O., and Veening, J.-W. (2012). Control of cell division in *Streptococcus pneumoniae* by the conserved Ser/Thr protein kinase StkP. Proc. Natl. Acad. Sci. U. S. A. 109, E905–913. 10.1073/pnas.1119172109.

43. Barendt, S.M., Land, A.D., Sham, L.-T., Ng, W.-L., Tsui, H.-C.T., Arnold, R.J., and Winkler, M.E. (2009). Influences of capsule on cell shape and chain formation of wild-type and *pcsB* mutants of serotype 2 *Streptococcus pneumoniae*. J. Bacteriol. 191, 3024–3040. 10.1128/JB.01505-08.

44. de Bakker, V., Liu, X., Bravo, A.M., and Veening, J.-W. (2022). CRISPRi-seq for genome-wide fitness quantification in bacteria. Nat. Protoc. 17, 252–281. 10.1038/s41596-021-00639-6.

45. Pinho, M.G., Kjos, M., and Veening, J.-W. (2013). How to get (a)round: mechanisms controlling growth and division of coccoid bacteria. Nat. Rev. Microbiol. 11, 601–614. 10.1038/nrmicro3088.

46. Briggs, N.S., Bruce, K.E., Naskar, S., Winkler, M.E., and Roper, D.I. (2021). The Pneumococcal Divisome: Dynamic Control of *Streptococcus pneumoniae* Cell Division. Front. Microbiol. 12, 737396. 10.3389/fmicb.2021.737396.

47. Balaban, N.Q., Helaine, S., Lewis, K., Ackermann, M., Aldridge, B., Andersson, D.I., Brynildsen, M.P., Bumann, D., Camilli, A., Collins, J.J., et al. (2019). Definitions and guidelines for research on antibiotic persistence. Nat. Rev. Microbiol. 17, 441–448. 10.1038/s41579-019-0196-3.

48. Kaldalu, N., Hauryliuk, V., and Tenson, T. (2016). Persisters-as elusive as ever. Appl. Microbiol. Biotechnol. 100, 6545–6553. 10.1007/s00253-016-7648-8.

49. Bui, N.K., Eberhardt, A., Vollmer, D., Kern, T., Bougault, C., Tomasz, A., Simorre, J.-P., and Vollmer, W. (2012). Isolation and analysis of cell wall components from *Streptococcus pneumoniae*. Anal. Biochem. 421, 657–666. 10.1016/j.ab.2011.11.026.

50. Bals, R., Wang, X., Zasloff, M., and Wilson, J.M. (1998). The peptide antibiotic LL-37/hCAP-18 is expressed in epithelia of the human lung where it has broad antimicrobial activity at the airway surface. Proc. Natl. Acad. Sci. 95, 9541–9546. 10.1073/pnas.95.16.9541.

51. LaRock, C.N., and Nizet, V. (2015). Cationic antimicrobial peptide resistance mechanisms of streptococcal pathogens. Biochim. Biophys. Acta 1848, 3047–3054. 10.1016/j.bbamem.2015.02.010.

52. Majchrzykiewicz, J.A., Kuipers, O.P., and Bijlsma, J.J.E. (2010). Generic and Specific Adaptive Responses of *Streptococcus pneumoniae* to Challenge with Three Distinct Antimicrobial Peptides, Bacitracin, LL-37, and Nisin. Antimicrob. Agents Chemother. 54, 440–451. 10.1128/aac.00769-09.

53. Bui, N.K., Turk, S., Buckenmaier, S., Stevenson-Jones, F., Zeuch, B., Gobec, S., and Vollmer, W. (2011). Development of screening assays and discovery of initial inhibitors of pneumococcal peptidoglycan deacetylase PgdA. Biochem. Pharmacol. 82, 43–52. 10.1016/j.bcp.2011.03.028.

54. Gonzalez, D.J., Campeau, A., and McGrosso, D.M. (2022). Protective vaccine antigen against streptococcal infection. Patent WO2022256310A1.

55. Cleverley, R.M., Rutter, Z.J., Rismondo, J., Corona, F., Tsui, H.-C.T., Alatawi, F.A., Daniel, R.A., Halbedel, S., Massidda, O., Winkler, M.E., et al. (2019). The cell cycle regulator GpsB functions as cytosolic adaptor for multiple cell wall enzymes. Nat. Commun. 10, 261. 10.1038/s41467-018-08056-2.

56. Stauberová, V., Kubeša, B., Joseph, M., Benedet, M., Furlan, B., Buriánková, K., Ulrych, A., Kupčík, R., Vomastek, T., Massidda, O., et al. (2024). GpsB Coordinates StkP Signaling as a PASTA Kinase Adaptor in *Streptococcus pneumoniae* Cell Division. J. Mol. Biol. 436, 168797. 10.1016/j.jmb.2024.168797.

57. Rismondo, J., Wamp, S., Aldridge, C., Vollmer, W., and Halbedel, S. (2018). Stimulation of PgdA-dependent peptidoglycan N-deacetylation by GpsB-PBP A1 in *Listeria monocytogenes*. Mol. Microbiol. 107, 472–487. 10.1111/mmi.13893.

58. Krawczyk-Balska, A., Korsak, D., and Popowska, M. (2014). The surface protein Lmo1941 with LysM domain influences cell wall structure and susceptibility of *Listeria monocytogenes* to cephalosporins. FEMS Microbiol. Lett. 357, 175–183. 10.1111/1574-6968.12518.

59. Kobayashi, K., Sudiarta, I.P., Kodama, T., Fukushima, T., Ara, K., Ozaki, K., and Sekiguchi, J. (2012). Identification and Characterization of a Novel Polysaccharide Deacetylase C (PdaC) from *Bacillus subtilis*. J. Biol. Chem. 287, 9765–9776. 10.1074/jbc.M111.329490.

60. Rahman, M.M., Zamakhaeva, S., Rush, J.S., Chaton, C.T., Kenner, C.W., Hla, Y.M., Tsui, H.-C.T., Uversky, V.N., Winkler, M.E., Korotkov, K.V., et al. (2024). O-glycosylation of intrinsically disordered regions regulates homeostasis of membrane proteins in streptococci. BioRxiv Prepr. Serv. Biol., 2024.05.05.592596. 10.1101/2024.05.05.592596.

61. Typas, A., Banzhaf, M., van den Berg van Saparoea, B., Verheul, J., Biboy, J., Nichols, R.J., Zietek, M., Beilharz, K., Kannenberg, K., von Rechenberg, M., et al. (2010). Regulation of peptidoglycan synthesis by outer-membrane proteins. Cell 143, 1097–1109. 10.1016/j.cell.2010.11.038.

62. Paradis-Bleau, C., Markovski, M., Uehara, T., Lupoli, T.J., Walker, S., Kahne, D.E., and Bernhardt, T.G. (2010). Lipoprotein cofactors located in the outer membrane activate bacterial cell wall polymerases. Cell 143, 1110–1120. 10.1016/j.cell.2010.11.037.

63. Midonet, C., Bisset, S., Shlosman, I., Cava, F., Rudner, D.Z., and Bernhardt, T.G. (2023). MacP bypass variants of *Streptococcus pneumoniae* PBP2a suggest a conserved mechanism for the activation of bifunctional cell wall synthases. mBio 14, e0239023. 10.1128/mbio.02390-23.

64. Paik, J., Kern, I., Lurz, R., and Hakenbeck, R. (1999). Mutational analysis of the *Streptococcus pneumoniae* bimodular class A penicillin-binding proteins. J. Bacteriol. 181, 3852–3856. 10.1128/JB.181.12.3852-3856.1999.

65. Domenech, A., Slager, J., and Veening, J.-W. (2018). Antibiotic-Induced Cell Chaining Triggers Pneumococcal Competence by Reshaping Quorum Sensing to Autocrine-Like Signaling. Cell Rep. 25, 2390–2400.e3. 10.1016/j.celrep.2018.11.007.

66. Lanie, J.A., Ng, W.-L., Kazmierczak, K.M., Andrzejewski, T.M., Davidsen, T.M., Wayne, K.J., Tettelin, H., Glass, J.I., and Winkler, M.E. (2007). Genome sequence of Avery’s virulent serotype 2 strain D39 of *Streptococcus pneumoniae* and comparison with that of unencapsulated laboratory strain R6. J. Bacteriol. 189, 38–51. 10.1128/JB.01148-06.

67. Mignolet, J., Fontaine, L., Sass, A., Nannan, C., Mahillon, J., Coenye, T., and Hols, P. (2018). Circuitry Rewiring Directly Couples Competence to Predation in the Gut Dweller *Streptococcus salivarius*. Cell Rep. 22, 1627–1638. 10.1016/j.celrep.2018.01.055.

68. Abramson, J., Adler, J., Dunger, J., Evans, R., Green, T., Pritzel, A., Ronneberger, O., Willmore, L., Ballard, A.J., Bambrick, J., et al. (2024). Accurate structure prediction of biomolecular interactions with AlphaFold 3. Nature 630, 493–500. 10.1038/s41586-024-07487-w.

69. Pettersen, E.F., Goddard, T.D., Huang, C.C., Meng, E.C., Couch, G.S., Croll, T.I., Morris, J.H., and Ferrin, T.E. (2021). UCSF ChimeraX: Structure visualization for researchers, educators, and developers. Protein Sci. Publ. Protein Soc. 30, 70–82. 10.1002/pro.3943.

70. de Jong, I.G., Beilharz, K., Kuipers, O.P., and Veening, J.-W. (2011). Live Cell Imaging of *Bacillus subtilis* and *Streptococcus pneumoniae* using Automated Time-lapse Microscopy. J. Vis. Exp. JoVE, 3145. 10.3791/3145.

71. Schindelin, J., Arganda-Carreras, I., Frise, E., Kaynig, V., Longair, M., Pietzsch, T., Preibisch, S., Rueden, C., Saalfeld, S., Schmid, B., et al. (2012). Fiji: an open-source platform for biological-image analysis. Nat. Methods 9, 676–682. 10.1038/nmeth.2019.

72. Sharma, V. (2018). ImageJ plugin HyperStackReg V5.6. Version v5.6 (Zenodo). 10.5281/zenodo.2252521 https://doi.org/10.5281/zenodo.2252521.

73. Dénéréaz, J., and Veening, J.-W. (2024). BactEXTRACT: an R Shiny app to quickly extract, plot and analyse bacterial growth and gene expression data. Access Microbiol. 6, 000742.v3. 10.1099/acmi.0.000742.v3.

74. Kulak, N.A., Pichler, G., Paron, I., Nagaraj, N., and Mann, M. (2014). Minimal, encapsulated proteomic-sample processing applied to copy-number estimation in eukaryotic cells. Nat. Methods 11, 319–324. 10.1038/nmeth.2834.

75. Käll, L., Storey, J.D., and Noble, W.S. (2008). Non-parametric estimation of posterior error probabilities associated with peptides identified by tandem mass spectrometry. Bioinforma. Oxf. Engl. 24, i42–48. 10.1093/bioinformatics/btn294.

76. Nesvizhskii, A.I., Keller, A., Kolker, E., and Aebersold, R. (2003). A statistical model for identifying proteins by tandem mass spectrometry. Anal. Chem. 75, 4646–4658. 10.1021/ac0341261.

77. Liu, X., Gallay, C., Kjos, M., Domenech, A., Slager, J., van Kessel, S.P., Knoops, K., Sorg, R.A., Zhang, J.-R., and Veening, J.-W. (2017). High-throughput CRISPRi phenotyping identifies new essential genes in *Streptococcus pneumoniae*. Mol. Syst. Biol. 13, 931. 10.15252/msb.20167449.

78. Bravo, A.M., Typas, A., and Veening, J.-W. (2022). 2FAST2Q: a general-purpose sequence search and counting program for FASTQ files. PeerJ 10, e14041. 10.7717/peerj.14041.

79. Deatherage, D.E., and Barrick, J.E. (2014). Identification of mutations in laboratory-evolved microbes from next-generation sequencing data using breseq. Methods Mol. Biol. Clifton NJ 1151, 165–188. 10.1007/978-1-4939-0554-6_12.

80. Clinical and Laboratory Standards Institute (CLSI). Performance Standards for Antimicrobial Susceptibility Testing. (2023). Clin. Lab. Stand. Inst. CLSI Perform. Stand. Antimicrob. Susceptibility Test. 33rd Ed CLS.

81. Hayashi, K. (1975). A rapid determination of sodium dodecyl sulfate with methylene blue. Anal. Biochem. 67, 503–506. 10.1016/0003-2697(75)90324-3.

82. White, R.M., Sessa, A., Burke, C., Bowman, T., LeBlanc, J., Ceol, C., Bourque, C., Dovey, M., Goessling, W., Burns, C.E., et al. (2008). Transparent adult zebrafish as a tool for in vivo transplantation analysis. Cell Stem Cell 2, 183–189. 10.1016/j.stem.2007.11.002.

83. Jim, K.K., Engelen-Lee, J., van der Sar, A.M., Bitter, W., Brouwer, M.C., van der Ende, A., Veening, J.-W., van de Beek, D., and Vandenbroucke-Grauls, C.M.J.E. (2016). Infection of zebrafish embryos with live fluorescent *Streptococcus pneumoniae* as a real-time pneumococcal meningitis model. J. Neuroinflammation 13, 188. 10.1186/s12974-016-0655-y.

