## Supplementary Figures for "A bacterial cell wall repair and modification system to resist host antibacterial factors"

### Supplementary information

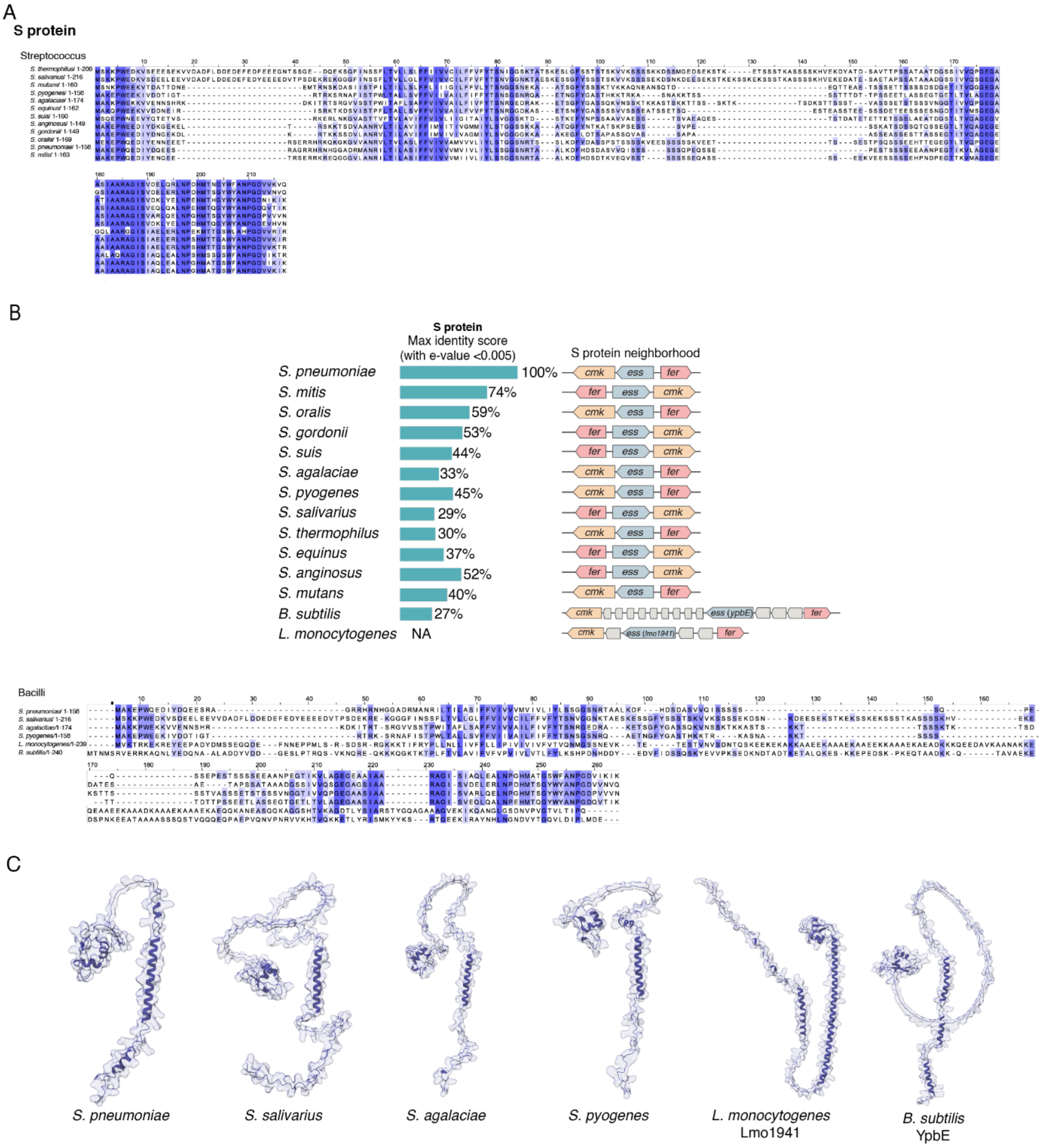


**Fig. S1**: **S protein conservation in Bacilli.** (**A**) Multiple sequence alignment of Streptococci S protein sequences. Sequence similarity are highlighted in dark blue. (**B**) Percentages indicate the percent identity for each species compared to *S. pneumoniae* S protein. *L. monocytogenes* S protein identity score compared to *S. pneumoniae* S protein is 21% using Clustal Omega. Gene co-occurrence in several bacilli genomes (data obtained from genome annotation in NCBI: see Methods). Multiple sequence alignment of possible S protein sequences of several bacilli. Sequence similarity are highlighted in dark blue. (**C**) AlphaFold model of the S protein of different bacilli bacteria.

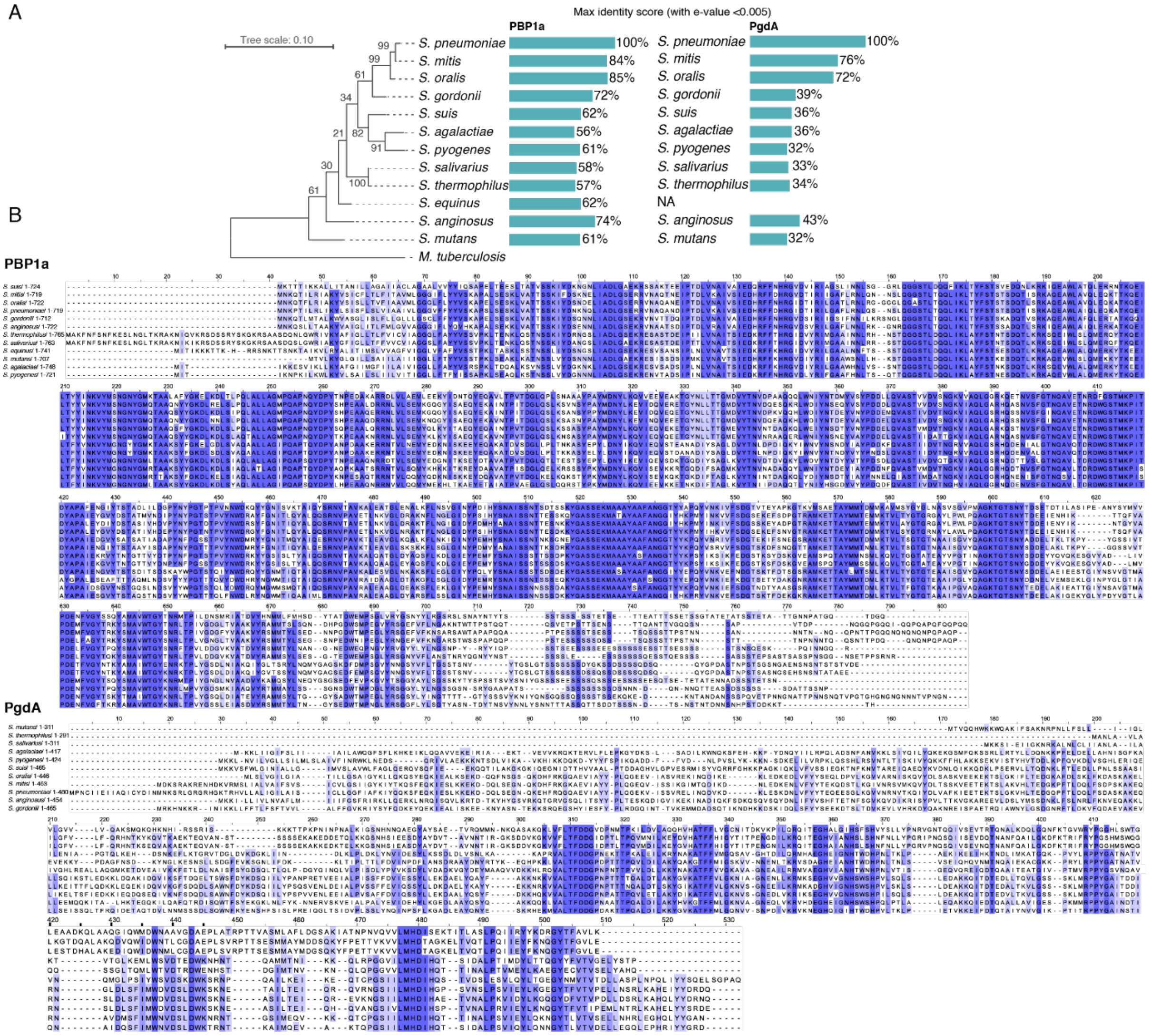


**Fig. S2**: (**A**) PBP1a and PgdA conservation in Streptococci. Lineage tree based on 16S rRNA sequence of each specie (see methods) was constructed using MEGA (Tamura et al., 2021). M*ycobacterium tuberculosis* was used as an outgroup and numbers represent bootstrap. Percentages indicate the percent identity of PBP1a and PgdA for each species compared to *S. pneumoniae* PBP1a and PgdA respectively. (**B**) Multiple sequence alignment of Streptococci PBP1a or PgdA sequences. Sequence similarity are highlighted in dark blue.

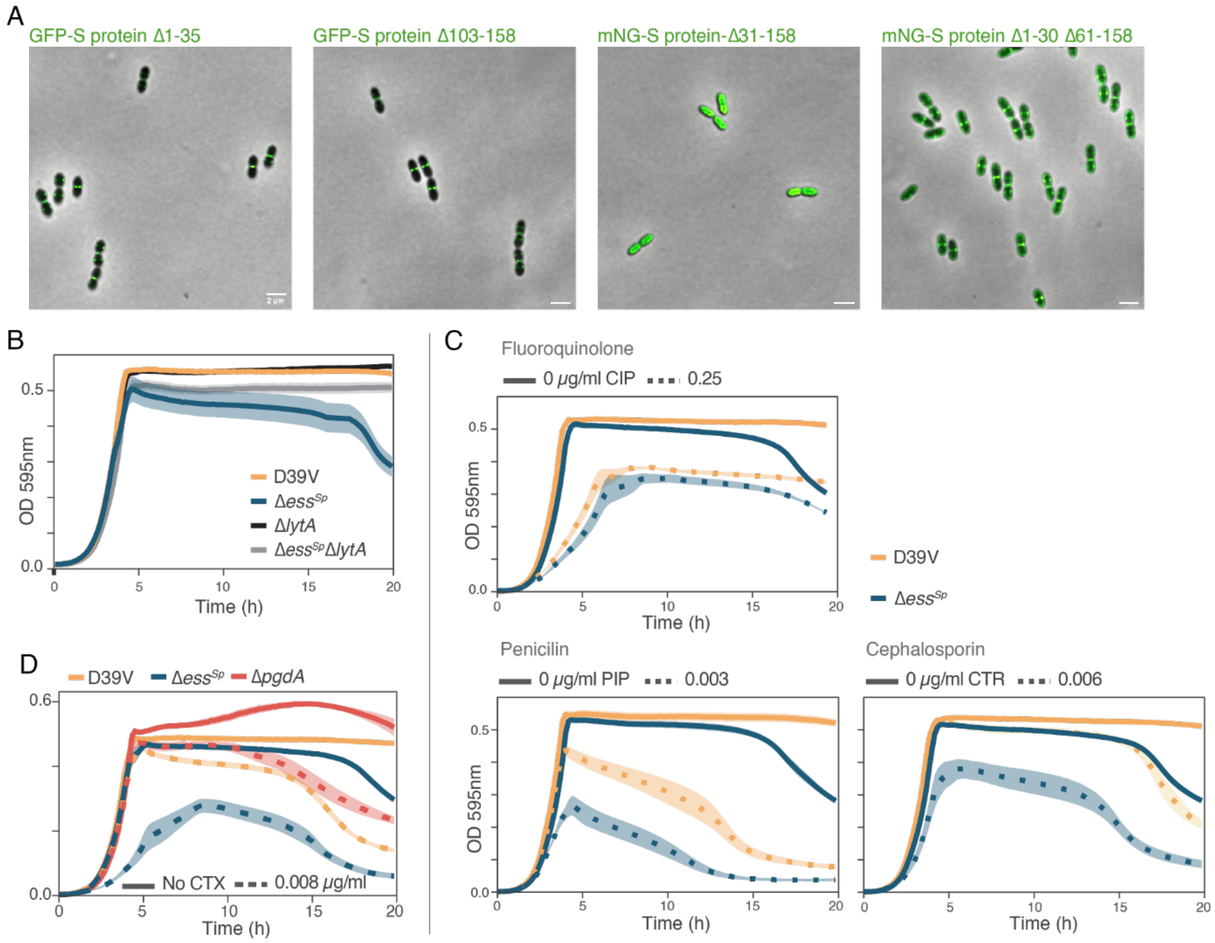


**Fig. S3**: (**A**) Deconvolved epifluorescence microscopy of live *S. pneumoniae* cells containing truncation fragments of S protein tagged with msfGFP (GFP) or mNeonGreen (mNG). The transmembrane (TM) domain is necessary for S protein septal localization. Scale bar = 2 μm. (**B**) Growth curve of pneumococcal cells deleted with either Δ*lytA* or Δ*ess*$^{Sp}$, or both at 37°C. Data are represented as the mean of n≥ 3 replicates and shading represents SEM. (**C**) Growth curves of WT or Δ*ess*$^{Sp}$ cells subject to diverse antibiotics; ciprofloxacin (CIP), piperacillin (PIP) or ceftriaxone (CTR) at 37°C. Δ*ess*$^{Sp}$ mutant is more susceptible to cell wall targeting antibiotics (PIP, CTR) than the wild type strain and has the same susceptibility to DNA targeting antibiotic (CIP). Data are represented as the mean of n≥ 3 replicates and shading represents SEM. (**D**) Growth curves of pneumococcal cells subject to cefotaxime (CTX) at 37°C show that the Δ*pgdA* mutants is not more susceptible compared to the wild type. Data are represented as the mean of n≥ 3 replicates and shading represents SEM.

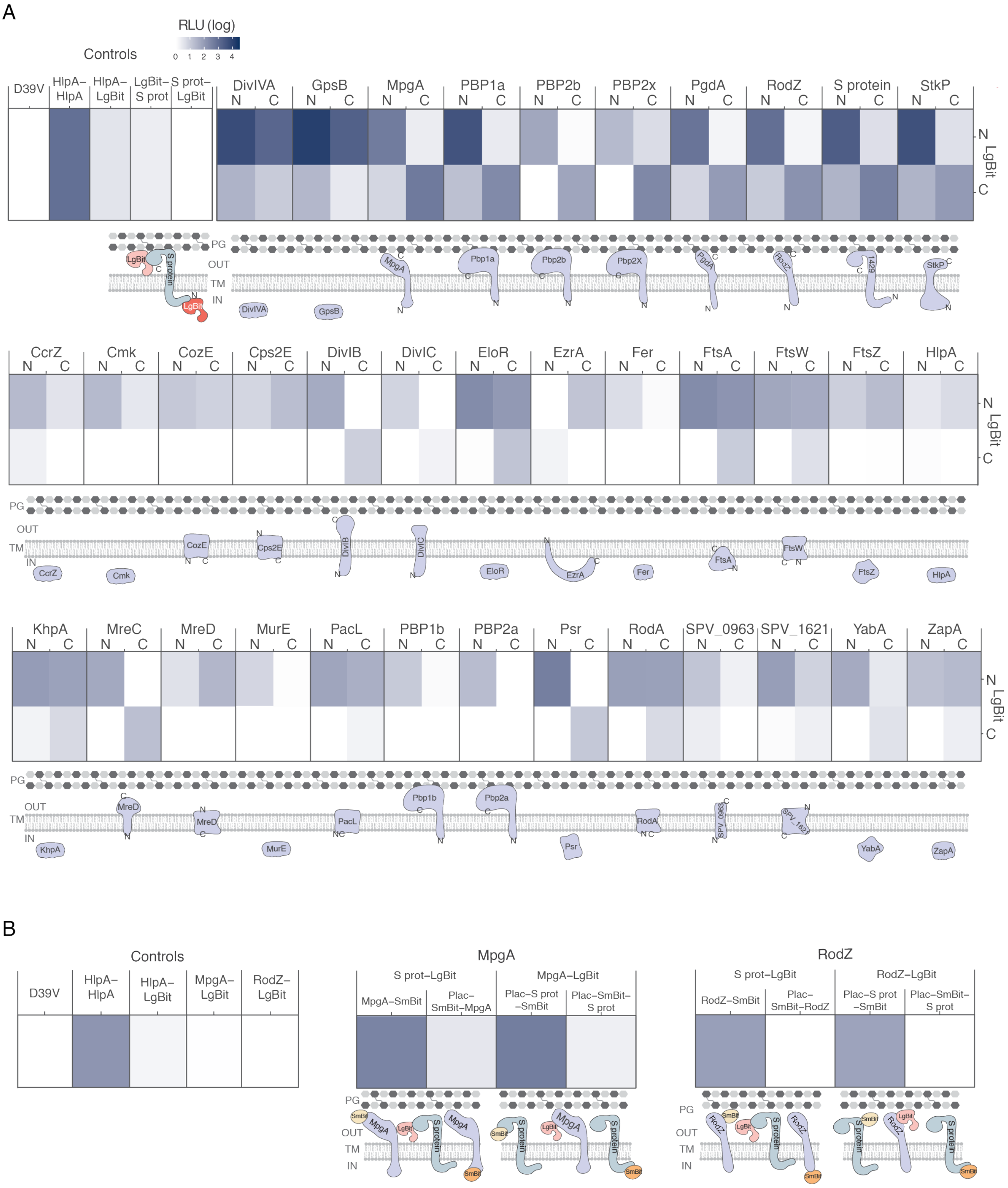


**Fig. S4: Split-luciferase assay.** (**A**) All proteins tested by split-luciferase assay. High relative luminescence unit (RLU) indicate close proximity (0 represent no proximity, 4 represent close proximity). HlpA-LgBit HlpA-SmBit (HlpA-HlpA) is used as a positive control. DivIVA, GpsB, MpgA, PBP1a, PgdA, RodZ, S protein and StkP are potential partners as they have high signal for two of the four combinations. In comparison, PBP2b and PBP2x have a weaker signal. Protein representations are not drawn to scale, and when relevant, C and N indicate the sub-cellular localization of the C- and N-terminal SmBit fusion respectively. (**B**) Fusion of the LgBit to the C-ter of MpgA and RodZ confirmed their close proximity to S protein.

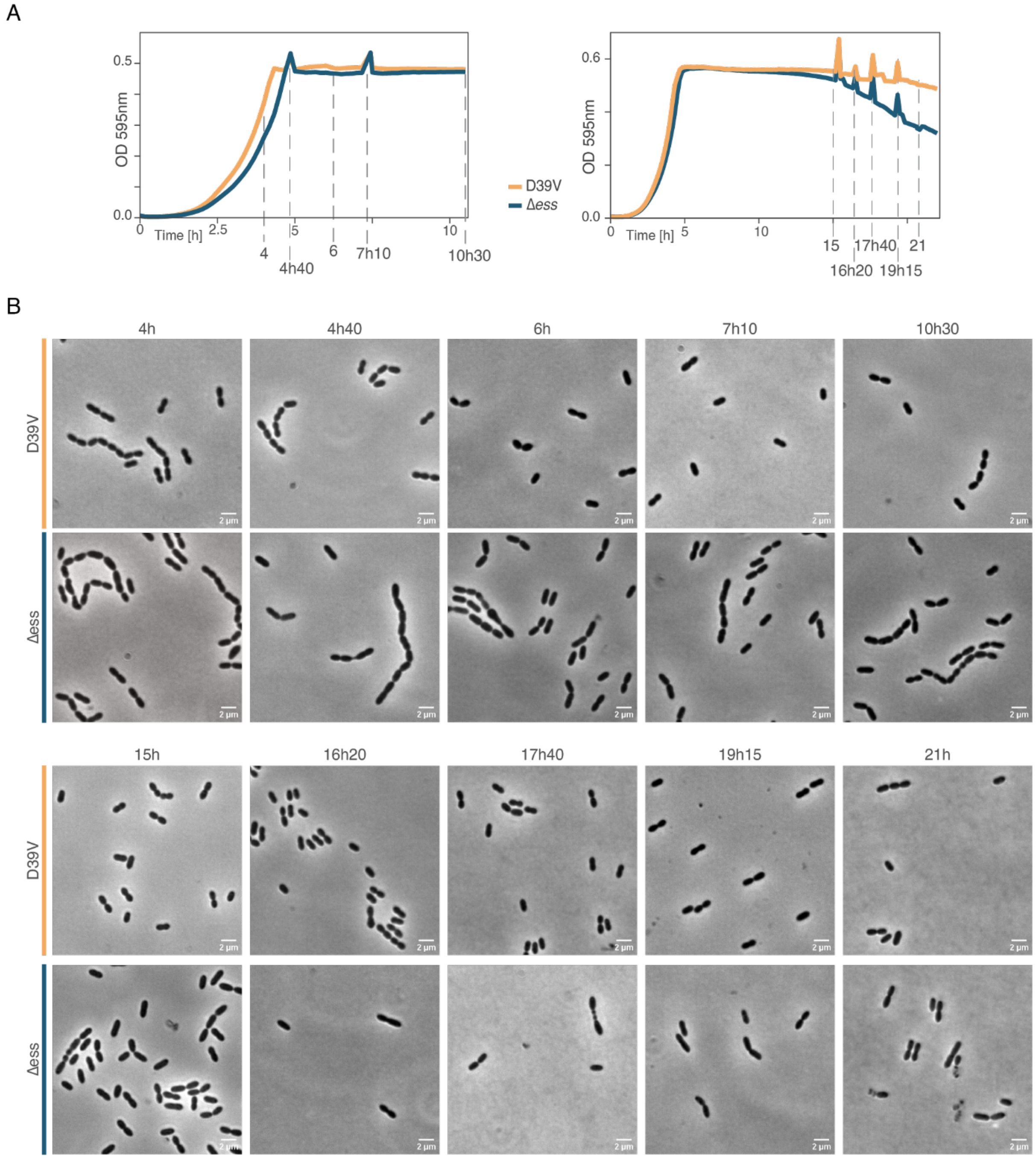


**Fig. S5: Time course experiment.** (**A**) Growth curves of Δ*ess*$^{Sp}$ deleted pneumococcal cells shows increased cell lysis compared to D39V wild type at 37°C. Dotted lines represent the times at which cells were harvested for microscopy. Peaks are due to the plate reader being opened to take samples for microscopy. (**B**) Phase contrast microscopy of Δ*ess*$^{Sp}$ deleted pneumococcal cells at different time point. Scale bar = 2 µm.

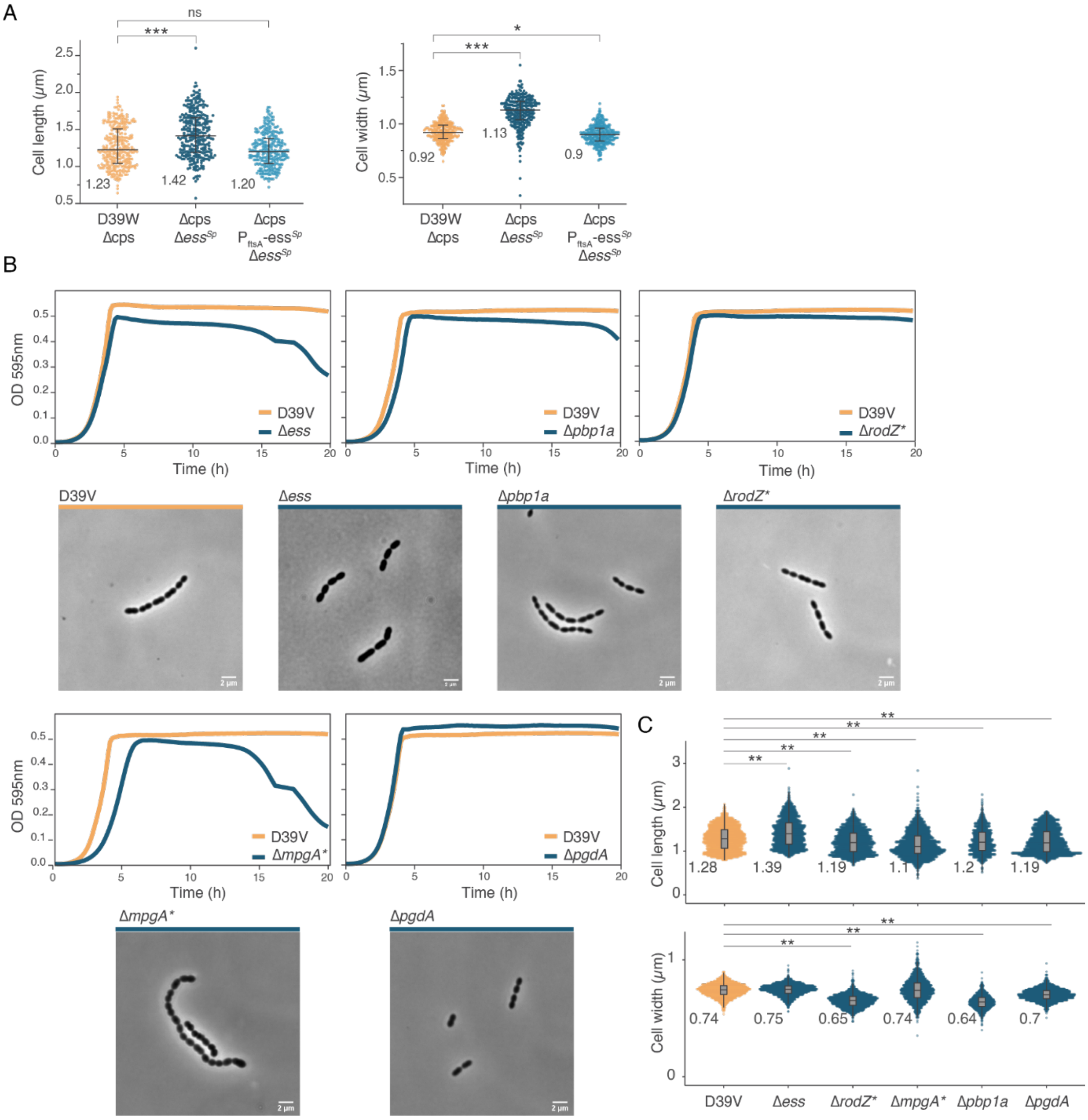


**Fig. S6: Phenotypes of single mutants without or with capsule.** (**A**) Quantitative analysis of phase contrast images show that Δ*cps* Δ*ess*$^{Sp}$ cells are significantly longer and wider than wild type D39W or complemented cells. 100 cells from each of 3 independent experiments were measured and plotted with scatter dot plot including median and interquartile bars. *, *** and ns denote $p<0.05$, $p<0.001$, not significant, respectively when compared to WT. Numbers represent the means. (**B**) Growth curve of the deletion of single gene (blue) compared to D39V wild type (yellow) at 37°C. Data are represented as the mean of n≥ 3 replicates. Phase contrast microscopy images of liquid culture of *S. pneumoniae* D39V wild type or upon single deletion of genes. Scale bar = 2 µm. Note that strains with an asterisk (Δ*rodZ**, for example) contain suppressor mutations described in Fig. 4B and Table S5. (**C**) Quantitative analysis of phase contrast images. The number of cells recorded ranges between 1000 and 2000 cells, numbers represent the means. ** $p<0.01$.

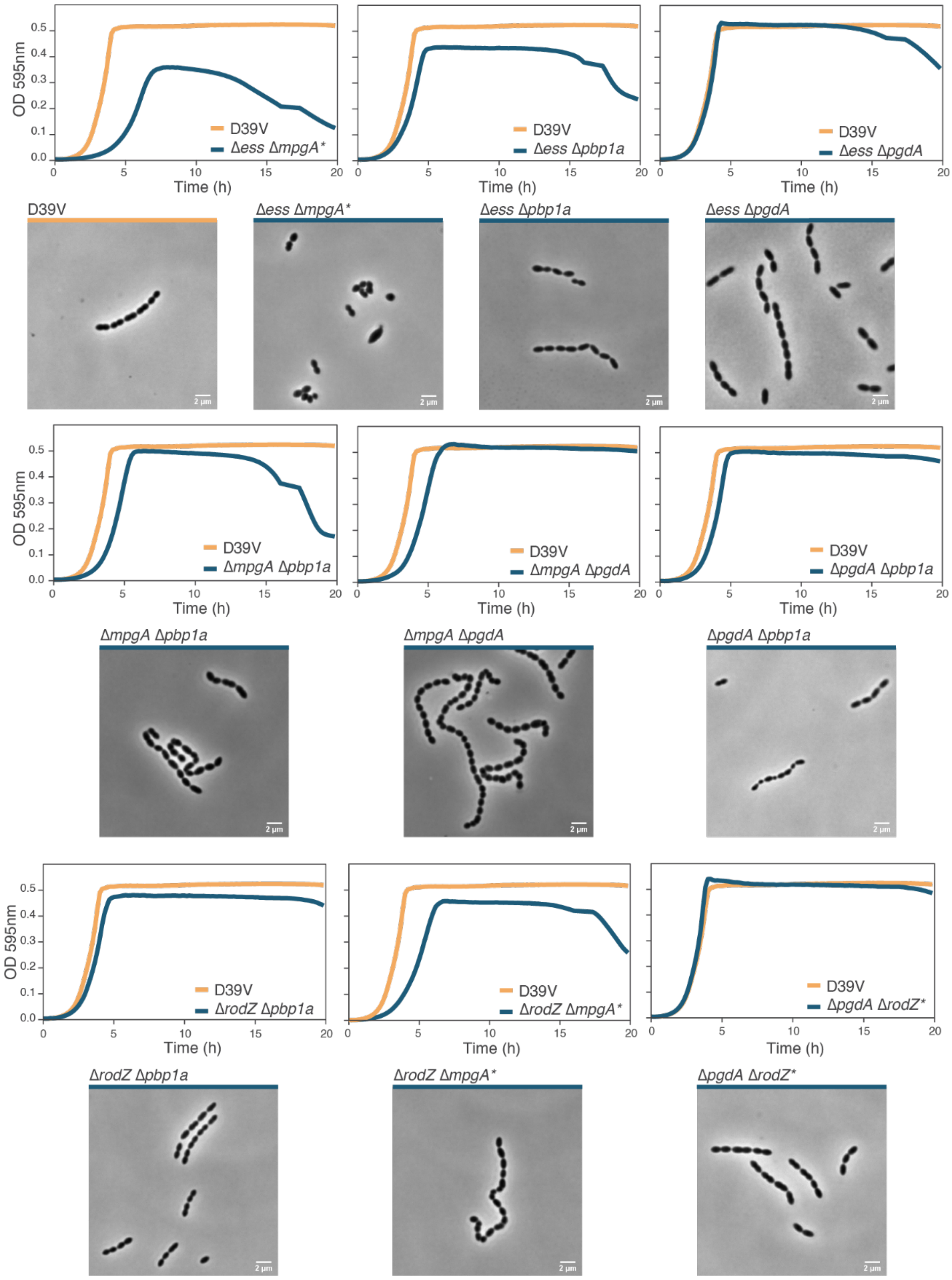


**Fig. S7: Double mutants phenotype.** Growth curve of the deletion of two genes (blue) compared to D39V wild type (yellow) at 37°C. Data are represented as the mean of n≥ 3 replicates. Phase contrast microscopy images of liquid culture of *S. pneumoniae* D39V wild type or upon double deletion of genes. Scale bar = 2 µm. Note that strains with an asterisk (Δ*ess* Δ*mpgA**, for example) contain suppressor mutations described in Fig. 4B and Table S5.

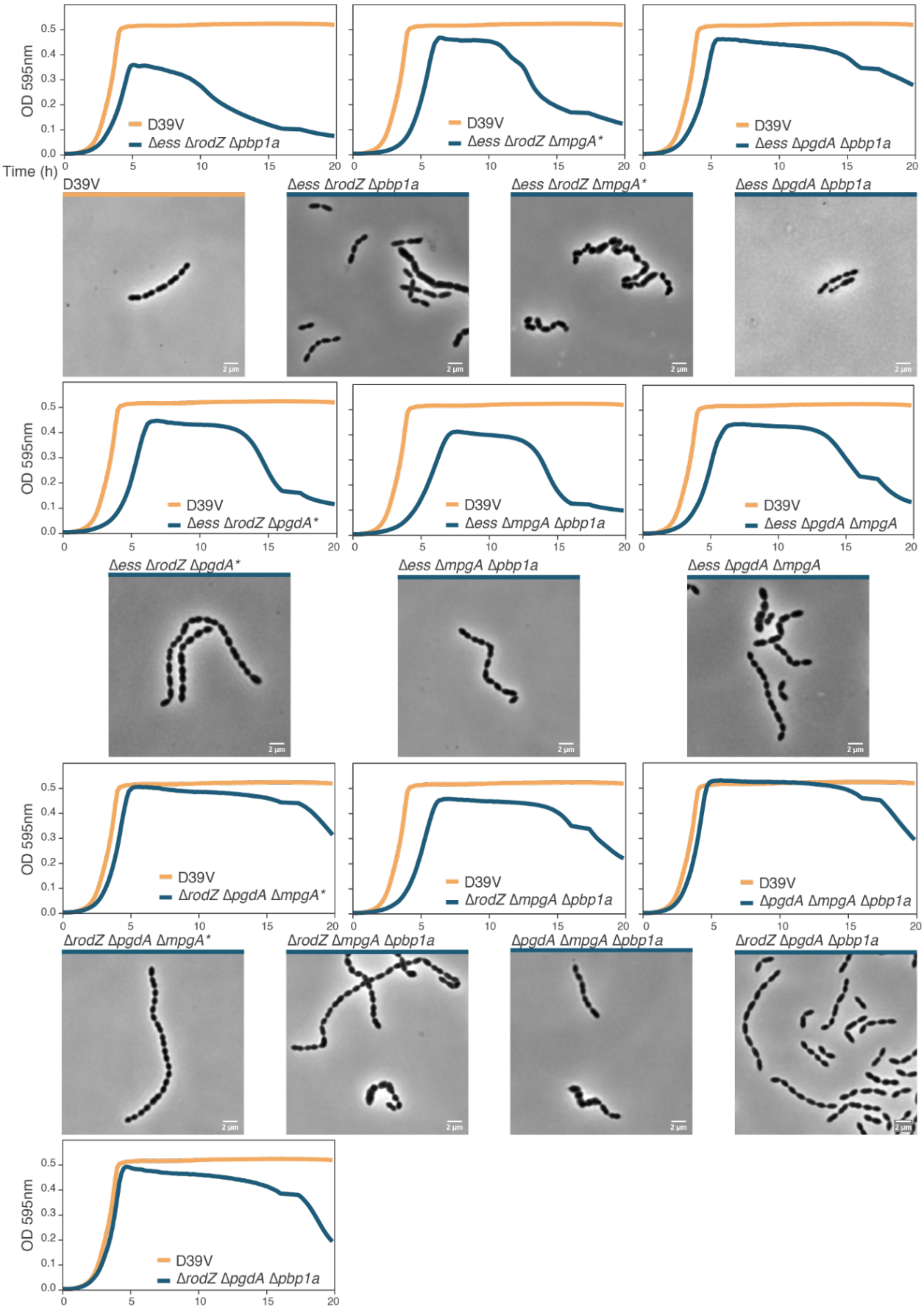


**Fig. S8: Triple mutants phenotype.** Growth curve of the deletion of three genes (blue) compared to D39V wild type (yellow) at 37°C. Data are represented as the mean of n≥ 3 replicates. Phase contrast microscopy images of liquid culture of *S. pneumoniae* D39V wild type or upon triple deletion of genes. Scale bar = 2 µm. Note that strains with an asterisk (Δ*ess* Δ*rodZ* Δ*mpgA**, for example) contain suppressor mutations described in Fig. 4B and Table S5.

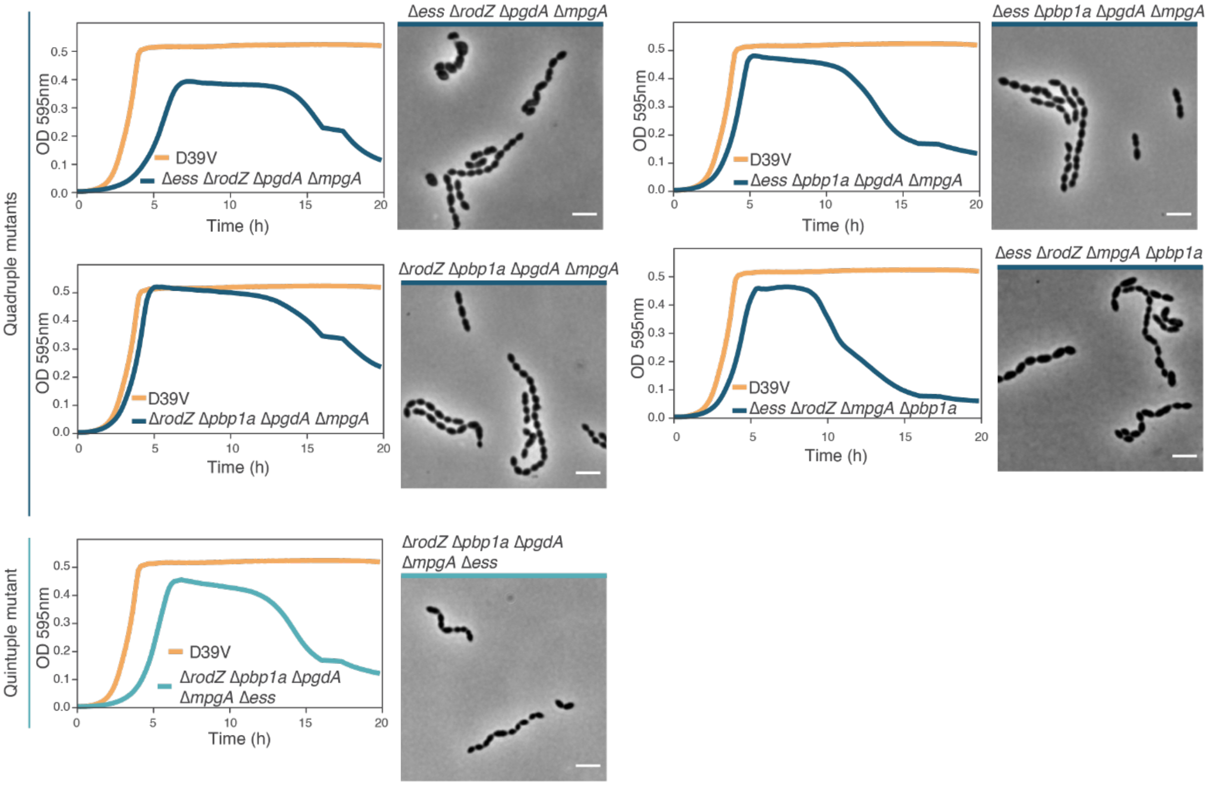


**Fig. S9: Quadruple and quintuple mutants phenotype.** Growth curve of the deletion of four (dark blue) or five genes (light blue) compared to D39V wild type (yellow) at 37°C. Data are represented as the mean of n≥ 3 replicates. Phase contrast microscopy images of liquid culture of *S. pneumoniae* D39V upon quadruple or quintuple deletion of genes. Scale bar = 2 µm.

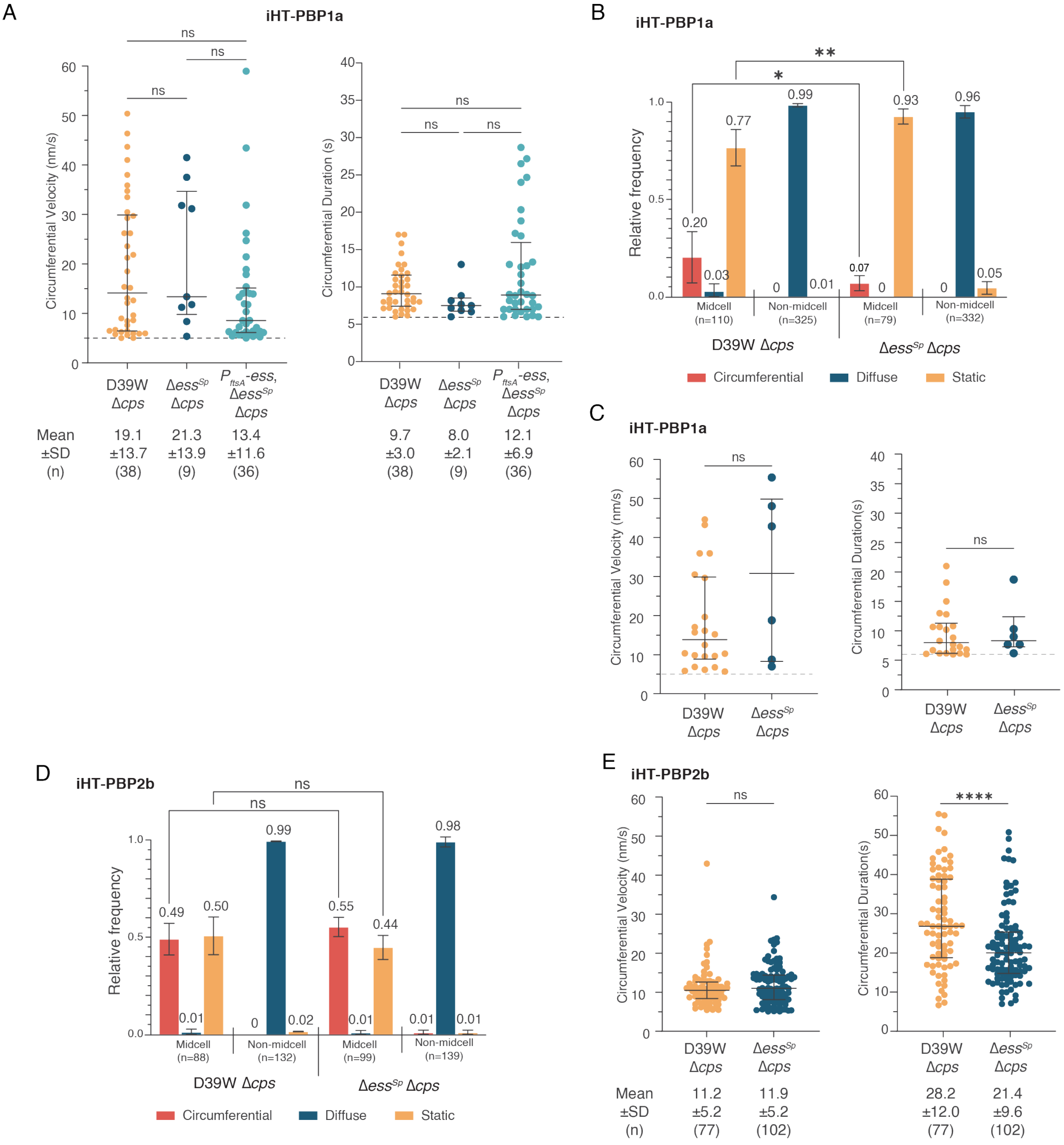


**Fig. S10:** (**A**) Deleting *ess* does not affect the velocity or duration of circumferentially moving iHT-PBP1a. All strains express a functional fusion of the HaloTag (iHT) domain fused to PBP1a. (left) Velocities and (right) durations of circumferentially moving HT-labeled single molecules of PBP1a are shown. Dots represent individual measurements, black horizontal line shows median, error bars denote interquartile range. Mean, standard deviation (±SD) and total number of circumferential molecules analyzed (*n*) from two biological replicates shown below graph. *ns* (not significant), $P > 0.05$. (**B-C**) Deleting *ess* decreases the frequency of circumferentially moving iHT-aPBP1a molecules, but does not affect their velocity. Both strains are merodiploids that express a functional fusion of the HaloTag (iHT) domain fused to PBP1a from the native locus of *pbp1a* as well as from an ectopic site under the control of a zinc-inducible promoter. (**B**) Movement patterns of HT-labeled single molecules of aPBP1a are shown. Bars represent the mean relative frequencies (±SD) of circumferential, diffusive and static molecules at midcell or away from the midcell (non-midcell) as determined by live cell TIRFm, with mean values shown above the bars. *n* = the total number of molecules analyzed from two biological replicates. * $P \leq 0.05$; ** $P \leq 0.01$; all other comparisons (not shown) were not significant, $P > 0.05$. (**C**) (left) Velocities and (right) durations of circumferentially moving HT-labeled single molecules of PBP1a are shown and plotted as described above. (**D-E**) Deleting *ess* does not affect the frequency or velocity of circumferentially moving molecules of iHT-PBP2b, but reduces their duration. Both strains are merodiploids with a construction similar as mentioned above with PBP2b instead of PBP1a. (**D**) Movement patterns of HT-labeled single molecules of PBP2b are shown. *n* = the total number of molecules analyzed from two biological replicates. *ns* (not significant), $P > 0.05$. (**E**) Velocities and durations of circumferentially moving HT-labeled single molecules of PBP2b are shown. Mean, standard deviation (±SD) and total number of circumferential molecules analyzed (*n*) from two biological replicates shown below graph. *ns* (not significant), $P > 0.05$; ****, $P < 0.0001$. See Methods (Single-molecule dynamics) for details of statistical tests.

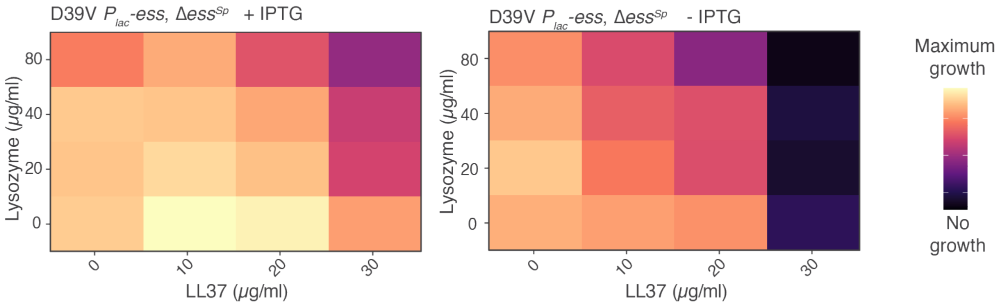


**Fig. S11: Pneumococcal susceptibility to host-induced damage in the absence of the S protein can be complemented.** Heatmap of the area under the curve (AUC) of the growth curves in liquid media. Lysozyme and LL-37 act synergistically on the Δ*ess*$^{Sp}$ mutant. Empirical AUC was plotted between 0-7 hours.

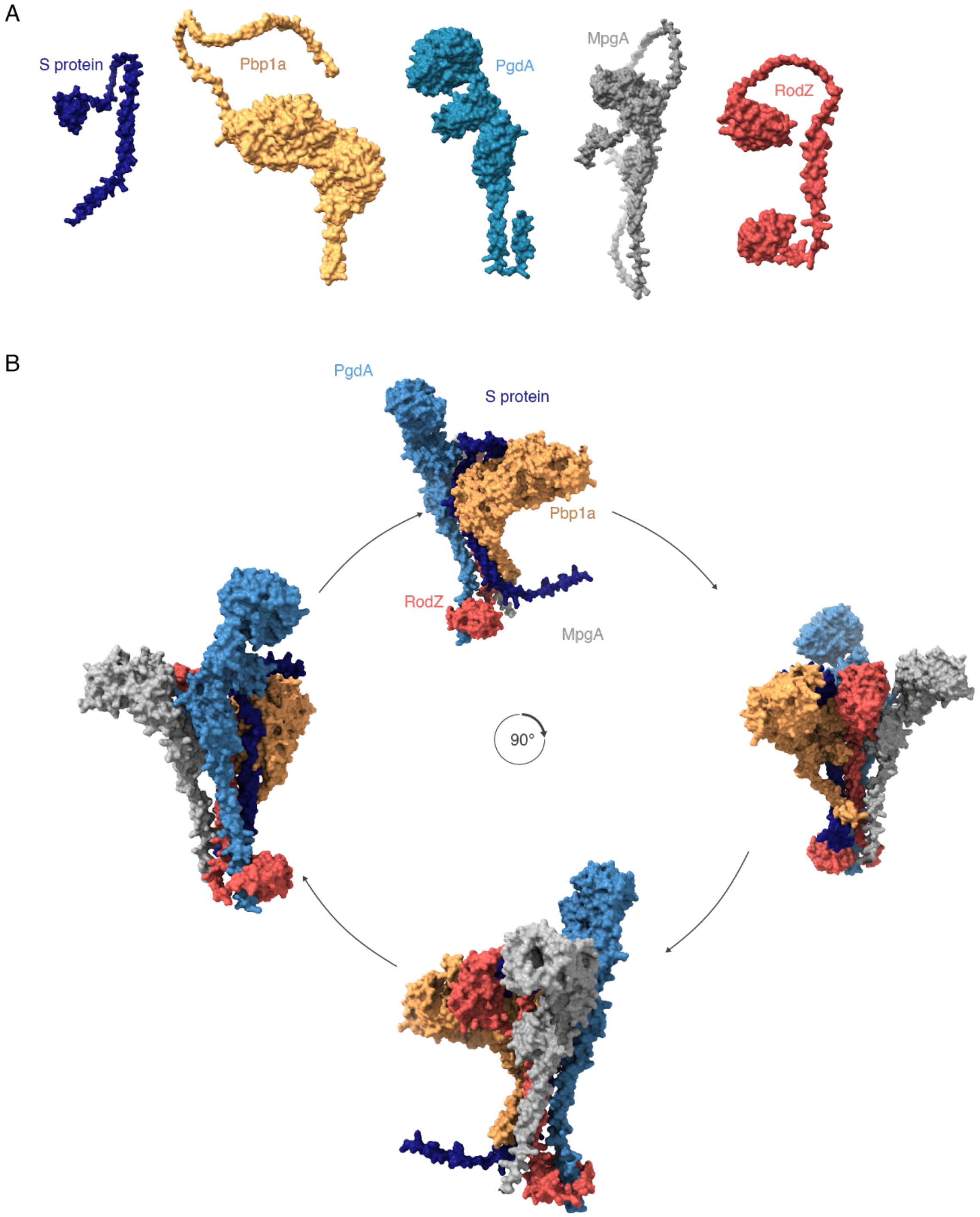


**Fig. S12:** (**A**) AlphaFold prediction of all five proteins generated separately using AlphaFold 3. (**B**) AlphaFold prediction of the potential complex formed by the S protein, PBP1a, PgdA, MpgA and RodZ. For clarity purposes, the amino-acids 651 to 719 of PBP1a and amino acid 1 to 173 of MpgA are not shown.

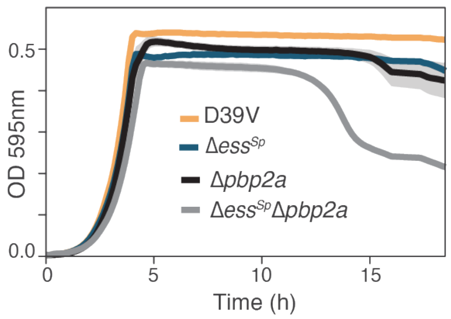


**Fig. S12:** Growth curve of pneumococcal cells deleted with either Δ*pbp2a* or Δ*ess*$^{Sp}$, or both at 37°C. Data are represented as the mean of n≥ 3 replicates and shading represents SEM.
